# Differential Expression Enrichment Tool (DEET): An interactive atlas of human differential gene expression

**DOI:** 10.1101/2022.08.29.505468

**Authors:** Dustin J. Sokolowski, Jedid Ahn, Lauren Erdman, Huayun Hou, Kai Ellis, Liangxi Wang, Anna Goldenberg, Michael D. Wilson

## Abstract

Differential gene expression analysis using RNA sequencing (RNA-seq) data is a standard approach for making biological discoveries. Ongoing large-scale efforts to process and normalize publicly available gene expression data enable rapid and systematic reanalysis. While several powerful tools systematically process RNA-seq data, enabling their reanalysis, few resources systematically recompute differentially expressed genes (DEGs) generated from individual studies. We developed a robust differential expression analysis pipeline to recompute 3162 human DEG lists from The Cancer Genome Atlas, Genotype-Tissue Expression Consortium, and 142 studies within the Sequence Read Archive. After measuring the accuracy of the recomputed DEG lists, we built the Differential Expression Enrichment Tool (DEET), which enables users to interact with the recomputed DEG lists. DEET, available through CRAN and RShiny, systematically queries which of the recomputed DEG lists share similar genes, pathways, and TF targets to their own gene lists. DEET identifies relevant studies based on shared results with the user’s gene lists, aiding in hypothesis generation and data-driven literature review.

**Highlights:** By curating metadata from uniformly processed human RNA-seq studies, we created a database of 3162 differential expression analyses.

These analyses include TCGA, GTEx, and 142 unique studies in SRA, involving 985 distinct experimental conditions.

The Differential Expression Enrichment Tool (DEET) allows users to systematically compare their gene lists to this database.

## INTRODUCTION

RNA sequencing (RNA-seq) is commonly used to measure genome-wide transcriptional abundance within and across biological samples (1). RNA-seq experiments typically compare RNA transcript abundances between two or more groups to calculate differentially expressed genes (DEGs) (2). Interpreting DEG results often involves gene ontology (GO) enrichment (3– 5), public gene co-expression network comparisons using a myriad of tools such GeneMANIA and EnrichR (6, 7), and directly comparing results to published studies. Currently, hundreds of thousands of human RNA-seq samples are publicly available within the Sequence Read Archive (SRA) (8, 9), and accessing these data efficiently and meaningfully is an important step in RNA-seq analysis.

Advances in large-scale analyses of publicly available RNA-seq data make it possible to interact with public data (5–7) systematically. Reprocessing publicly available RNA-seq data before interpreting their results is essential because of technical variation inherent to the experimental and analytical steps of an RNA-seq study. Large-scale projects like *recount*, toil-recompute, and ARCHS4 (10–13), remove unwanted variation in analysis by developing and applying efficient computational strategies to consistently align and enumerate RNA-seq data across tens of thousands of samples simultaneously (10–13). Briefly, *recount2* stores consistently reprocessed RNA-seq data from ∼70,000 samples. Of these samples, ∼20,000 of them originated from consortia with complete metadata, with 9,538 samples from The Cancer Genome Atlas (TCGA) (14, 15) and 11,284 samples from the Genotype-Tissue Expression Consortium (GTEx) (16). Modern public consortia of re-reprocessed RNA-seq data now combine to store over a million human and mouse RNA-seq samples (10, 12). Using *recount*, ARCHS4, and other consistently processed RNA-seq databases, researchers can download and compare RNA-seq samples to their data (17, 18) without worrying about any technical heterogeneity in RNA-seq data analysis.

High-quality metadata is fundamental to analyzing consistently processed RNA-seq data properly. However, metadata is not always consistently stored. For example, Ellis *et al*., 2018 analyzed the metadata of 49,564 human RNA-seq samples stored with the Sequence Read Archive (SRA) and found that sex was only reported in 3640 (7.3%) of those samples (19). The Gene Expression Omnibus (GEO) and ArrayExpress (20, 21) provide guidelines and greatly facilitate the submission of RNA-seq data and associated metadata. MetaSRA also improved the organization of public metadata by developing a semi-automated metadata normalization process to convert published metadata to a format comparable to metadata stored in the Encyclopedia of DNA Elements (ENCODE) (22, 23). While these efforts and others facilitate the pairing of RNA-seq studies with metadata, there are still considerable inconsistencies in metadata between datasets regarding metadata organization and sample missingness. One solution to the problem of incomplete metadata was addressed by Ellis et al., 2018 using the PhenoPredict (19) package to improve the metadata within *recount2* (11). Specifically, PhenoPredict (19) trained a metadata classifier from TCGA and GTEx RNA-seq data stored within *recount2* before annotating the remaining ∼50,000 SRA samples within *recount2*, resulting in uniform metadata across *recount2*. Additional projects like recount-brain use a third party to manually annotate a consistent set of metadata for brain RNA-seq samples within *recount (24)*. Together, consistent RNA-seq count data and metadata allow for the development of pipelines to conduct high-throughput differential expression analysis.

Several existing robust methodologies allow for querying extensive systematically processed RNA-seq data. For example, Enrichr and the Expression Atlas from ArrayExpress allow for the systematic querying of gene lists (7, 25). Enrichr included co-expression from consistently reprocessed RNA-seq data in ARCHS4 (12, 25), while GenomicSuperSignatures applies a principal component analysis approach to 536 studies to study-associated gene-expression patterns (26). Two other tools have also reprocessed DEG lists to aid in biological discovery. Specifically, The Expression Atlas (25, 27) contains co-expression and DEGs from many species and experiments, including over 330 pairwise DE comparisons from human RNA-seq alone. Secondly, Crow et al., 2019 used 635 pairwise human DE comparisons from consistently processed microarray data from Gemma database to better understand common distributions of DEGs (28, 29). These methods highlight the value of uniformly processed RNA-seq data, metadata, and differential expression; however, there is still a considerable need for larger-scale atlases of interactive DE.

In this study, we describe the Differential Expression Enrichment Tool (DEET), a database and bioinformatic package that allows users to query systematically generated differential gene expression results from published RNA-seq studies. DEETs database contains 3,162 consistently processed human pairwise differential gene expression comparisons from studies within *recount2 (11)*, spanning 99 tissues, 55 cell lines, and 985 conditions (486 from SRA, 433 from TCGA, 66 from GTEx). DEET allows users to input a list of genes with relevant coefficients (e.g., p-value, fold-change, GWAS effect size) to systematically query the gene expression and pathway enrichment profiles of thousands of consistent gene lists through gene set enrichment and correlational analyses. DEET and its database can be accessed via a freely-available library of DE comparisons, R package (https://cran.rstudio.com/web/packages/DEET/index.html), and Shiny App (https://wilsonlab-sickkids-uoft.shinyapps.io/DEET-shiny/).

## MATERIALS AND METHODS

The purpose of the Differential Expression Enrichment Tool (DEET) is to facilitate comparing user-defined lists of differentially expressed genes (DEGs) against a uniformly computed and annotated compendium of DEGs (Figure 1A-B). To build the DEET database, we computed a compendium of 3162 unique, consistently processed human DEG comparisons, and developed supporting software (R package and Shiny app) to interact with the DEG compendium. For each pairwise comparison, DEGs were identified using a custom pipeline that uses factor analysis of metadata and DESeq2 for differential analysis. Next, for each DEGs list, DEET performs GO term and TF target enrichment analysis. The pre-computed DEGs and enrichment results are stored in DEET.

**Figure 1.**
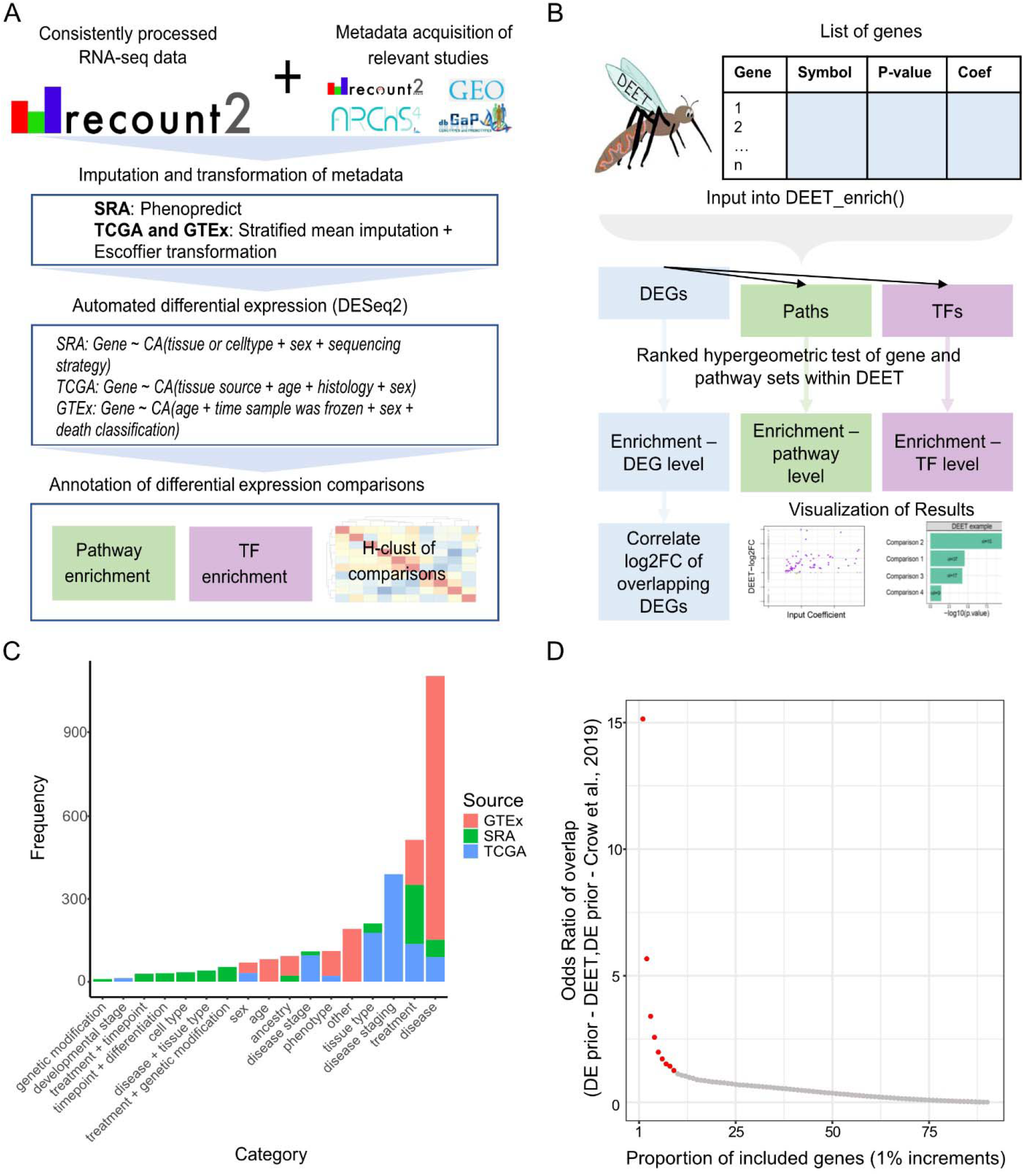
Overview of the Differential Expression Enrichment Tool (DEET). A) Schematic of how the consistently processed DEGs were computed and annotated. B) Flowchart of DEET’s primary analysis. C) Barplot of the number of comparisons from each DEG-comparison category in DEET. Categories plotted were derived from the categories labeling 635 pairwise DE comparisons from Microarray studies in the Gemma database (Crow *et al*., 2019). We added sex, developmental staging, and combinations of treatments as additional categories. Bars are coloured by source (i.e., GTEx, TCGA, and SRA). D) Scatterplot showing the odds ratio of overlapping common DEGs between the DEET database and Crow et al., 2019. The X-axis represents the proportion of included genes, ranked from most common to least common. For example, the “1%” point includes genes in the top 1% most common in either DEET or Crow et al., 2019. The Y-axis represents the odds ratio of over-representation of shared genes at each increment. Points in red represent increments with a significant over-representation of shared DEGs between the DEET database and Crow et al., 2019.

### Data acquisition

All RNA-seq count data were acquired from the “recount” R package using the “download_study” function with default parameters (11). Metadata from studies with the SRA, TCGA, and GTEx were acquired from multiple sources.

#### SRA

Metadata for studies within SRA was acquired by using the “all_metadata” function in the “recount” R package and supplemented with the “human_matrix_v9.h5” file in ArchS4 (8, 11, 12). Samples stored within recount-brain (24) was further supplemented with “add_metadata(source = “recount_brain_v2”)” using the “recount” R package (11). Specifically, we extracted overlaid sample metadata in recount2 and ArchS4 by their “geo_accession”. We then added the “title” variable from Archs4 to the metadata stored in recount2 (11, 12, 19) (Supplementary File S1). Lastly, we downloaded brief descriptions of each study from the DRA compendium (https://trace.ddbj.nig.ac.jp/DRASearch/).

#### TCGA

Metadata for The Cancer Genome Atlas (TCGA) was acquired from the “recount” R package using the “all_metadata” function (11, 15).

#### GTEx

Publicly available metadata for the Genotype-Tissue Expression (GTEx) consortium was acquired with the “all_metadata” function (11, 16). Privately available metadata for GTEx was acquired using dbGap (phs000424.v9) with all required ethical approvals and data protection.

### Metadata pre-processing

We needed to streamline the metadata with SRA, GTEx, and TCGA before we could perform differential analysis within each study and tissue. Streamlined metadata in combination with consistently reprocessed RNA-seq count data allowed for high-throughput differential expression analysis within each sample source.

#### SRA

Metadata across different studies submitted to SRA is inherently inconsistent. Accordingly, within SRA, we focused on metadata compatible with the PhenoPredict R package (19). These compatible metadata variables are tissue, cell type, sample source, sex, and sequencing strategy. Specifically, if the authors reported values for these variables, then PhenoPredict converted the consistent variable names across datasets (e.g., “reported tissue”, and they are populated with the reported value. PhenoPredict also matches reported metadata with predicted metadata variables based on the RNA-seq profile of each sample trained on the metadata and RNA-seq profiles within GTEx (19). For the DEET database, we used the author’s reported metadata value and the predicted metadata if the reported metadata was unavailable.

#### TCGA

Metadata was first manually processed to remove possible inconsistencies. Specifically, we manually adjusted and merged drug names based on spelling errors and generic and brand names, respectively (e.g., ibuprofen vs. Advil). Variables where values contained different units (e.g., body temperature measured in Celsius vs. Fahrenheit) were also corrected so that every value adhered to the most common unit. For example, if the majority of body temperatures were reported in Celsius, then every sample changed their reported body temperature to Celsius. Missingness of continuous variables was populated with a mean imputation stratified by sex. Missingness of categorical variables was populated with an “unknown” label.

#### GTEx

We did not detect metadata requiring manual corrections within GTEx. Like TCGA, missing continuous variables were populated with a mean imputation stratified by sex, and missing categorical variables were populated with an “unknown” label.

### Comparison exclusion and inclusion criteria for the DEET database

Several criteria needed to be met for a comparison to be included in the DEET database. These inclusion and exclusion criteria were consistent across TCGA, SRA, and GTEx. Comparisons were filtered if they had fewer than three biological replicates in each condition, if conditions were generic identifications (e.g., Patient ID 1-5 vs. 6-10), if conditions were compared across different tissues, or if the comparison had a complete stratification of metadata (e.g., all “drug-control” were female and all “drug-treated” were male). Time-series and stepwise dosage comparisons kept the original reference point to each timepoint and stepwise timepoints while non-linear timepoints (e.g., Time-2 vs. Time-4 if Time-3 was present) were filtered.

Comparisons where one condition was “NA” or “unknown” were filtered for interpretability. Lastly, studies with more than three comparisons were filtered so that each treatment was only compared to an untreated control (e.g., each TCGA-drug was compared to the untreated condition). We removed these “treatment-a vs. treatment-b” comparisons to avoid having DEET be primarily populated with permutations of DE comparisons that are challenging to interpret.

Studies with three comparisons (i.e., Control, Treatment 1, and Treatment 2) include every pairwise difference, including Treatment 1 vs. Treatment 2, as it only added one extra comparison. Lastly, after DE was performed (See “High-throughput differential expression analysis”), comparisons with more than 10,000 DEGs and fewer than 5 DEGs were filtered.

While the exclusion criterion for comparisons originating from TCGA, SRA, and GTEx was the same, the inclusion criteria for comparisons originating from these sources differed

#### SRA

SRA is a repository of unique studies. Therefore, comparison variables across studies in SRA were inconsistent. Comparisons were included in DEET if they passed the general exclusion criterion. The remaining comparisons were then paired with the description of each study found within the DRASearch (https://ddbj.nig.ac.jp/search). Comparisons reflecting the study description are included, and comparisons that do not reflect the study description are flagged and only included if the comparison was not confounded by the primary comparison.

Five of these studies, SRP043162 (30), SRP063978 (31), SRP063980 (31), SRP064561 (32), SRP067214 (33), and SRP050892 (34), each had multiple timepoints and tissues with two conditions (98 comparisons total). Our high-throughput pipelines would have treated these features as blocking factors instead of variables to stratify pairwise comparisons. Accordingly, we completed these DE comparisons manually and provided them with the “SRA-manual” source identification. Lastly, 14 studies (21 comparisons) had their metadata manually supplemented with the recount-brain (24) dataset. Metadata from recount-brain (24) did not influence how DEs were calculated, but it did influence how these DE comparisons were described.

#### TCGA

Over 10,000 samples contained sample information mapped to the same metadata table. Accordingly, variables from this curated TCGA metadata table were manually selected for their potential to provide biologically meaningful comparisons. Specifically, we included variables describing the tumour such as tumour presence, reoccurrence, stage, grade, histological diagnosis, and subdivision. In addition, we included variables describing tumour treatment (e.g., follow-up, drug treatment, and surgery performed). Variables specific to individual cancer types (e.g., Estrogen receptor positivity, KRAS mutation presence) were included and automatically filtered from irrelevant cancers due to DEET’s database exclusion criterion because other tumour types contained missing or unknown cancers. In addition, we included ordinally annotated medical conditions (e.g., presence of diabetes, presence of heart disease, chronic pancreatitis). Lastly, population-level variables, namely sex and weight, were included. Weight was compared using body mass index (BMI) and was grouped into broad categories provided by the Centre for Disease Control and Prevention (https://www.cdc.gov/healthyweight/assessing/bmi/adult_bmi/index.html).

#### GTEx

Like in TCGA, metadata variables within GTEx were chosen from GTEx’s library of clinical variables (https://www.ncbi.nlm.nih.gov/projects/gap/cgi-bin/GetListOfAllObjects.cgi?study_id=phs000424.v4.p1&object_type=variable) for their ability to yield interpretable DE. Firstly, all clinical ordinal variables (e.g., presence of pneumonia at the time of death -yes or no) were included. Population variables, namely sex, age, race, BMI (with the same criteria as in TCGA), and Hardy Scale (i.e., death circumstances), were also included. Ages were binned into 20-year periods (i.e., 20-39, 40-59, and 60-79 years).

### High-throughput differential expression analysis

Each pairwise comparison has a different number of samples, different sample stratification, and potentially different combinations of categorical and continuous metadata to control for. Furthermore, each major source of samples contained a different set of metadata to control for (Figure 1A). Accordingly, our high-throughput differential analysis pipelines needed to be flexible for different experimental designs and variability in metadata. We used the variables below to account for population-level metadata within SRA, TCGA, and GTEx.

#### SRA

We control for tissue or cell type, sequence strategy, and sex.

#### TCGA

We control for tissue source, age, histological subtype, and sex.

#### GTEx

We control for age, time passed until sample freezing, Hardy Scale, and sex.

If we are measuring DEGs in a variable that we typically control for, for example, sex differences, then we do not control for sex in that comparison. We accounted for the variability in experimental designs within the DEET database by applying automated correspondence analysis to each comparison. Specifically, continuous metadata (e.g., age, sample freezing time, etc.) underwent an Escoffier transformation using the “ours” R package (35, 36). Categorical metadata (e.g., sex, tissue, etc.) underwent a disjunctive transformation using the “ours” R package. We then reduced these metadata into a smaller set of explanatory variables with a correspondence analysis (CA) using the “epCA” function in ExPosition (35). We used a correspondence analysis instead of a principal component analysis (PCA) because the metadata used to control for comparisons within DEET were mixed (i.e., continuous, and categorical).

Then, we generated a Screeplot of the eigenvectors for every CA and every comparison. We picked the number of components using the elbow of each graph. Together, every pairwise comparison controlled for variables appropriate to the variation in their metadata.

Differential gene expression analysis for all pairwise comparisons was completed with the likelihood ratio test in the DESeq2 R package (2). The appropriate number of factors as measured by the CA were used to reduce DE. Genes were considered differentially expressed if they had an FDR-adjusted p-value < 0.05. The “downregulated” group was decided alphabetically, as not all comparisons had a clear “case” and “control”.

Next, we performed pathway enrichment for every pairwise DE comparison. Specifically, we inputted all genes in each comparison into ActivePathways (37). We selected genes detected in each comparison as the statistical background. Genes with an FDR-adjusted p-value <0.05 were labeled significant. Enriched pathways included both up-regulated and down-regulated genes.

For both pathway and TF enrichment, we used an FDR-adjusted p-value to correct for enriched pathways. All pathways, regardless of significance, were returned. Pathways were derived from a homogeneous gene-set database (http://download.baderlab.org/EM_Genesets/). Specifically, we used the “Human_GO_AllPathways_with_GO_iea_June_01_2021_symbo.gmt” pathway enrichment file, where we included paths with 15-2000 genes. Transcription factor (TF) targets were derived from the “Human_TranscriptionFactors_MSigdb_June_01_2021_symbol.gmt” TF target file, where we included TFs with 15-5000 genes.

### Display of the differential expression comparisons within the DEET database

Every comparison was named using the following format: *Study ID*: *Cell/Tissue type*. condition *1* vs *condition 2*. When available, cell and tissue types were identified from internal metadata and the study summaries from the DRA compendium otherwise. In addition, for every pairwise comparison, the study name, source (SRA, TCGA, GTEx, and SRA-manual) (8, 15, 16), description from the DRA compendium, the number of samples (total, up-condition, and down-condition), samples (total, up-condition, down-condition), tissue (including tumour from TCGA), number of DEs (total, up-condition, down-condition), age (mean +-sd), sex, top 15 DEGs - up, top 15 DEGs - down, top 5 enriched pathways, and top 5 enriched TFs (Supplementary File S1) are provided. PubMed IDs are also available for studies selected from SRA. Lastly, each pairwise comparison was given an overall category based on a DE category list decided in Crow *et al*., 2019 (29). We also added the additional categories of sex, age, and combinations of categories (e.g., treatment + timepoint) to accommodate our additional comparisons.

### Comparing DE comparisons in the DEET database against their original studies

We compared a subset of the pairwise DE comparisons we recomputed against the same comparisons stored within the supplemental data of their original studies.

#### SRA

DEGs from Lin41 treatments vs. control in ESCs were obtained from Supplementary file S2 from Worringer *et al*., 2014 (38). DEGs from timepoints after FOXM1 inhibition MCF-7 cells were obtained from the GEO file (GSE58626) associated with Gormally *et al*., 2014 (39).

#### TCGA

Sex differences in DEGs for twelve different cancers summarized in Table 1 of Yaun *et al*., 2016 (40) were obtained by contacting the authors directly.

#### GTEx

Sex differences in DEGs for seventeen tissues were obtained from Supplementary Table 3 from Lopez-Ramos *et al*., 2020 (41), For all comparisons in the DEET database and the original study, we applied an absolute-value fold change cutoff of 1.5 and an FDR-adjusted p-value < 0.05 cutoff. For each matching comparison, we evaluated the over-representation of overlapping DEGs between the original study and DEET’s evaluation of DEGs with a Fisher’s Exact-test of overlapping genes using “cellmarker_enrich” in the scMappR R package (42, 43), where the background is the number of genes detected in the original study. We then tested whether applying the DEET enrichment tool to each original DE comparison would enrich for their analogous DEG comparison within the DEET database. We outputted the rank that the analogous comparison was enriched. If the analogous comparison was not ranked one (i.e., the most enriched study), we outputted whether every more strongly enriched comparison contained the same primary variable (e.g., sex differences in a different tissue). Then, we measured the log2(Fold-change) similarity of genes that overlapped between the two studies using a Pearson’s correlation. P-values for these Fisher’s-exact tests and correlations were FDR corrected, and the log2(Fold-change) of the genes designated as DE in the original study or the DEET database were plotted using the “ggplot2” R package (44).

### Implementation

We provide an R package, DEET, and Shiny applet that allows users to query a list of their genes against our 3162 consistently computed DEG lists. The DEET R package, can be installed from CRAN (Supplementary File S2; page 1), and the Shiny applet can be found at (https://wilsonlab-sickkids-uoft.shinyapps.io/DEET-shiny/). We also provide a workflow for users to query and visualize their DEGs against the DEET database (Supplementary File S1). We only query significant DEGs in the DEET R package and Shiny App. Both data sets can be downloaded with the DEET_data_download() function in the R package. The DEGs from each pairwise comparison within the DEET database are also stored in the gene-matrix transpose (*.gmt) format, allowing users to incorporate the DEET database with other pathway enrichment tools such as g:Profiler and GSEA (3, 4).

The primary function of the DEET R package is to allow users to query their list of DEGs against the consistently computed DEGs within the DEET database by using the function DEET_enrich(). The optimal input into DEET’s enrichment function, DEET_enrich(), is a data frame of genes (human gene symbols) with an associated p-value and coefficient (e.g., Fold-change) in conjunction with a list of genes designating the statistical background. First, DEET internally applies ActivePathways (37) function to the user’s gene list to identify enriched GO’s and TF’s using the same GO and TF datasets stored within the DEET database. DEET then uses ActivePathways (37) again to compute the enrichment of DEET comparisons at the gene, GO, and TF levels. ActivePathways (37) used all detected genes as the statistical background, Brown’s p-value fusion method, and an FDR-adjusted p-value cutoff of 0.05. Then, DEET_enrich() enriches the users’ inputted genes, pathways, and TF targets against the DEET database’s DEGs, pathways, and TF targets. Then, DEET_enrich() computes the Spearman’s and Pearson’s correlation between the coefficients of the user’s gene list and the log2(Fold-change) of DEGs within enriched pairwise comparisons in the DEET database. Finally, the p-values of these correlations are corrected with an FDR correction. Together, DEET’s enrichment tool returns significantly enriched studies based on overlapping DEGs, pathways, and TFs.

Optionally, DEET_enrich() may be used with a generic gene list (i.e., without P-values or coefficients). We assume an inputted list is unordered or in decreasing order of significance. If the gene list is ordered, we evenly space their p-value, with the least significant p-value being 0.049. The Pearson’s correlation between the inputted gene list and the DEGs within the DEET database is excluded. If the inputted gene list is unordered, then all p-values are set to 0.049, and both Spearman’s and Pearson’s correlations between the users’ inputted genes and the DEGs within the DEET database are excluded. If users do not provide a background set of genes, we assume the background set is all genes detected within the DEET database.

The DEET R package also contains plotting functions to summarize the most significant studies based on each enrichment test and correlation within DEET_enrich(). The process_and_plot_DEET_enrich() function plots barplots of the most enriched studies based on gene set enrichment (ActivePathways (37)) of the studies enriched studies based on overlapping DEGs, pathways, and TF targets. DEET also generates scatterplots of the most enriched studies based on Spearman’s correlation analysis. All plots are generated using ggplot2 (44), and DEET_enrich() returns the ggplot2 (44) objects for each plot to allow researchers to customize plots further.

Lastly, the DEET R package contains a function called DEET_feature_extract(), allowing researchers to identify genes whose log2Fold-change is associated with a response variable (e.g., fold-change of the gene of interest, and whether the study investigates cancer, etc.). Genes are extracted by calculating the coefficients from a Gaussian family elastic net regression using the “glmnet” R package (45, 46) and Spearman’s correlation between every gene and the response variable. If the response variable is categorical (e.g., comparison category), features are extracted by calculating the coefficients from a multinomial family elastic net regression and an ANOVA (47) between each category within the response variable. Lastly, if the response variable is ordinal (e.g., enriches for TNFa pathway yes/no, Cancer study yes/no, etc.), features are extracted using a binomial family elastic net regression and a Wilcoxon’s test (48) between the two categories within the response variable.

### Clustering of studies within the DEET database

Pairwise correlation analysis was completed within every study in the DEET database. Specifically, we took genes DE in at least one of the studies for each pair of studies and completed a Pearson’s correlation of their FDR-adjusted p-values. The R^2^ of these pairwise correlations were populated into a correlation matrix. We then computed the Euclidean distance matrix of the absolute value of the correlation matrix before performing a hierarchical clustering correlation matrix using the Ward.D (49) method and with a height cut-off of 100. The correlation matrix was clustered and plotted with the Pheatmap R package (44, 49). Median proportions of overlapping DEGs within each cluster were calculated by making a comparison-by-comparison matrix and populating it with the number of intersecting genes. Then, each row of the matrix was divided by the number of DEs in that row’s comparison. The median of this matrix was then calculated and represented by the barplot. For example, a value of 0.075 for cluster 5 means that “on average, a comparison within cluster 5 will share 7.5% of their DEGs with another comparison within cluster 5”. Finally, we annotated the biological and hallmark gene-sets for each cluster using ActivePathways, using Brown’s p-value fusion method.

### Case Study: Evaluating TNFa response in human endothelial cells

We acquired the full edgeR results of differential expression analysis in both the intronic RNA-seq and exonic RNA-seq from the original authors of Alizada *et al*., 2021 (50, 51). All detected genes in Alizada *et al*., 2021 (51) were used as the statistical background. Genes were separated into up-regulated and down-regulated based on false-discovery rate using the authors’ cut-offs of FDR < 0.1 and absolute-value log2(Fold-change) of 0.6. Then, each gene list was inputted into the DEET_enrich() function using default parameters. We also generated matrices of all FDR-adjusted p-values where each row is a gene, and each column is an RNA-seq type (i.e., intronic RNA-seq and exonic RNA-seq). Genes with a log2(Fold-change) > 0 had their FDR set to 1 to focus on downregulated genes. These matrices were inputted into ActivePathways (37) using default parameters. The *gmt file inputted into ActivePathways was the full list of DEGs stored within the DEET database and can be accessed with DEET_data_download().

## RESULTS

### Summary of the Differential Expression Enrichment Tool: Atlas and R package

The Differential Expression Enrichment Tool (DEET) facilitates hypothesis generation and provides biological insight from user-defined differential gene expression results. To use DEET, users input a list of genes with an associated p-value and summary statistic (i.e., fold-change). DEET performs GO term- and TF target-enrichment analysis using this gene list. DEET compares a) the gene list itself against a database of the precomputed DEGs within this study and b) enriched GO terms and potential regulatory TFs with precomputed enrichment results. DEET returns a set of RNA-seq experiments with similar results together with the genes and pathways responsible for the overlap between studies. Finally, DEET provides functions to visualize and report enrichments.

DEET interacts with a consistent set of 3162 human DE analyses that we calculated. Specifically, the total of 3162 comparisons were selected based on sample numbers and the interpretability of the comparisons. In total, 405 studies in *recount2*, the reprocessed RNA-seq count data used to recompute these DEG sets, contained at least five samples and one variable with two or more groups. After study filtering, 142 of these 405 studies remained to recompute differential analysis. Specifically, 162 studies were filtered due to insufficient sample size in one group and/or improper dispersions in DESeq2. The remaining 98 studies were filtered because their metadata variables with multiple conditions did not meet the DEET databases inclusion criteria (see “Materials and Methods” for details). Briefly, these criteria included study-relatedness, metadata stratification, confounding, studies containing bulk or cell-sorted RNA-seq rather than single-cell biosamples, and interpretability of comparisons. Additionally, only potentially meaningful DE comparisons from within the original study and tissue (e.g., TCGA samples were not compared to GTEx samples, liver samples were not compared to kidney samples) were included (Figure 1A). It is important to note that filtered studies were not necessarily intended for differential analysis, and there was not an inherent flaw in the original studies but an incompatibility with DEET. Lastly, while no entire study was filtered because of the number of DEGs, 246 comparisons were filtered for containing more than 10,000 or fewer than 5 DEGs.

Comparisons in GTEx (N = 1594 comparisons) (16) and TCGA (N = 957 comparisons) (15) were chosen based on whether the metadata had discrete options in their clinical metadata sheets. The primary variable comparisons from SRA (N = 611 comparisons across 142 studies) (8) were chosen based on their relationship to the author’s reported study description, which we added to DEET’s metadata. To provide an overview of the 985 types of DE comparisons in the DEET database, we sorted comparisons into 26 combinations of DE categories originally defined by Crow et al., 2019 (29), with most categories related to “disease” or “treatment” (Figure 1C).

DEET uses a ranked hypergeometric test provided by ActivePathways to compare user-provided gene list to pre-computed DEGs, (37). Unlike the gene sets stored within GO and pathway databases, the gene lists used by DEET are weighted by p-value and fold-change. DEET correlates the DEG coefficients with the fold-changes of a user’s DEG list and tests if other studies are changing in a similar pattern. Lastly, DEET uses enriched GOs and TFs based on the user’s gene list to identify studies with similar pathway enrichments using the hypergeometric test in ActivePathways (37). Lastly, DEET provides software for data visualization of enriched gene lists.

### Global patterns of differentially expressed genes within the DEET database

We first investigated the number of samples within each comparison within the DEET database. Specifically, we found a median of 127, 141, and 12 samples per comparison from TCGA, GTEx, and SRA sources, respectively. After accounting for the ratio of samples in each condition (see “Materials and Methods” for details), there was a “scaled” sample size of 26, 13, and 7. As expected, we found that the number of DEGs was positively correlated with the ratio-scaled number of samples in every source (Supplementary Figure S1). Furthermore, when accounting for the ratio in sample size, the variance in the total number of DEGs also decreases as the sample size increases (Supplementary Figure S2).

Previously, Crow *et al*., 2019 used 635 pairwise human DE comparisons from consistently processed microarray data from the Gemma database (28, 29). To develop a “DE prior” statistic, a multifunctionality analysis optimizing the rank (52) of common DEGs that were predictive of gene expression in most studies was used (29). Their DE prior highlighted that genes related to sex, cellular response, extracellular matrix, and inflammation were commonly DE regardless of comparison, while housekeeping genes were uncommonly DE. Furthermore, due to the unbiased nature of the DE comparisons used to predict their DE prior, they predicted these DEGs to be robust across consortia. Therefore, we generated a DE prior for the DEET database to be able to compare whether the overall patterns of differential expression within the DEET database replicate those in Crow *et al*., 2019.

We found that building a DE prior from the DEGs stored within the DEET database yielded a correlated ranking of DEGs (p-value = 2.64 × 10^−171^, rho = 0.215) to the DE prior in Crow *et al*., 2019. Furthermore, the top 1% of DE genes in each “DE prior” list were significantly overlapping (FDR-adjusted p-value = 1.37 × 10^−18^, OR = 15.3), with 26 overlapping genes primarily related to the Y chromosome and inflammation (Figure 1D, Supplementary Figure S3). We then repeated this analysis at 1% intervals. We found that the top 10% of genes significantly overlapped between the “DE prior” from Crow *et al*., 2019 (29) and the DE prior from the DEET database (Figure 1D, Supplementary Figure S3). Together, the global patterns of DEG frequency within the DEET database replicate established differential expression patterns.

### Distribution of DEG comparisons and pathways within the DEET database

After profiling the DEGs within the DEET database, we investigated how the 3162 comparisons clustered based on their DE profile. We expected comparisons to be clustered by shared underlying biology and experimental design; however, many comparisons originate from population-level comparisons in large consortium datasets (e.g., age, sex, time of death, presence of pneumonia, etc., in GTEx). Accordingly, population versus experimental RNA-seq designs, such as those found in SRA, may also drive cluster structure. We indeed found that the comparison source played a substantial role in cluster formation, with 7/23 clusters composed entirely from GTEx comparisons and 1/23 clusters composed exclusively of TCGA comparisons (Supplementary Figure S4A-B, Supplementary File S1). While TCGA is a population-level cohort, much of the metadata stored within TCGA is related to specific treatments (i.e., drug treatment). Like the sample source, the tissue of origin within the DE comparison also contributed to cluster identification. For example, clusters 20 and 24 were composed almost exclusively of GTEx comparisons in EBV-transformed lymphocytes, and clusters 22 and 23 contained almost exclusively GTEX comparisons in different brain regions (Supplementary Figure 4B).

We investigated how many DEGs overlap between all pairwise comparisons within a cluster. We found that clusters primarily annotated by shared experimental design (i.e., clusters 1, 3, 4, 5, 6, 7, and 16) shared an average of 22.0% (4.8%-44.4%) of their DEGs with another comparison within the same cluster (Supplementary Figure 4C). In contrast, clusters defined by source (TCGA, GTEx, and SRA) or tissue only shared 7.1% (3.9%-14.2%) of their DEGs with another comparison within the same cluster, which is significantly less (one-tailed Student’s t-test, p = 0.035) (Supplementary Figure 4C). Using ActivePathways (37) which allows for data fusion of p-values merging across different DE comparisons before conducting gene set enrichment, we annotated each cluster with GO (Supplementary Figure 4D) and the 50 Hallmark gene sets (Supplementary Figure 4E). Many clusters contained enrichment for development and immune response pathways in the Hallmark and the GO gene sets. For example, the “Humoral immune response” gene ontology was in the top 5 most enriched pathways for 7/23 clusters (Supplementary Figure 4D), and the “Inflammatory response” was in the top 5 most enriched Hallmarks in 12/23 clusters (Supplementary Figure 4E). In addition, the “Kras signaling -down” hallmark gene set was in the top 5 most enriched gene sets in 21/23 clusters (Supplementary Figure 4E). This strong and consistent enrichment of KRAS signaling likely reflects a bias towards cancer-related experiments in the DEET database. Specifically, there are 957 comparisons from TCGA, and all considered at least cancer-related, 47 comparisons in GTEx investigating cancer, and 134 comparisons in SRA where “cancer” or “tumour” were part of the DE comparison name or description.

### Differential expressed genes within the DEET database reflect the findings in the original studies

We next evaluated how the gene lists within the DEET database reflect the DEGs reported in the original studies. We chose publicly available comparisons from each primary source within the DEET database (GTEx, TCGA, and SRA). To verify if our DE comparisons made from GTEx data correspond to previously published analyses, we compared the pairwise analysis of sex differences within 17 tissues to what was reported in the original study (Lopez-Ramos *et al*., 2020) (41). To verify our DE analysis of TCGA data, we compared our results for the pairwise sex differences within the 12 tumour types to what was reported in the original study (Yuan *et al*., 2016) (40). To verify our comparisons in SRA, we chose two studies: DEGs measured from a) MCF-7 cells after FOXM1 inhibition (control t=0 vs. 3, 6, and 9 hours) (Gormally *et al*., 2014) (39) and b) Lin41-1 knockdown, and Lin41-2 knockdown in human embryonic stem cells (Worringer *et al*., 2014) (38). As expected, we found that each DEG list obtained from the original study either enriched for its own comparison as the single most enriched gene list (6/6 comparisons from SRA, 4/12 comparisons from TCGA, 12/17 comparisons from GTEx) or enriched for a study within the same source and comparison type but in a different tissue. For example, sex differences in glioblastoma multiforme (GBM) stored within the supplementary files of Yuan *et al*., 2016 enriched for DEET-computed sex differences in Glioblastoma (GBM), the fifth most significant comparison, while the most significantly enriched comparison was sex differences in Uveal melanoma (UVM) within the TCGA cohort (15) (Supplementary Table S1). We also found that every pairwise comparison from these studies had a highly significant overlap in DEGs and highly correlated fold-changes in overlapping DEGs (Supplementary Table S1, Supplementary Figure S5). We captured 31.4%-87.1% of the original DEGs, which is in line with differences that can occur when comparing any two commonly used differential analysis approaches to the same RNA-seq count matrix (53).

Lastly, when looking at the total number of DEGs, we found a similar number or, in most cases, more DEGs between all the comparisons within the DEET database compared to the original studies (Supplementary Table S1). Differences in alignment, gene counting and normalization, and differential analysis all influence gene DEG detections and dispersions, thus impacting the total number of DEGs. In particular, DEET-specific non-coding DEG detection partially explains why DEET detects more DEGs than many of the original comparisons.

Specifically, DEET-specific DEGs are, on average, 6.8x (0.65-36.3) more likely to be non-coding genes than DEGs shared between the DEET database and the original study (Supplementary Figure S6). Overall, the automated differential pipeline DEET used to calculate DEGs accurately captured the DEGs from their original studies.

### DEET identifies relevant studies when applied to TNFa-mediated inflammation

To demonstrate how DEET can be used to explore user-generated DEG lists, we took our lab’s previously published analysis of human aortic endothelial cells (HAoEC) treated with proinflammatory cytokine tumour necrosis factor-alpha (TNF) (51). TNFa stimulation activates the transcription factor complex NF-κB and drives rapid proinflammatory gene expression. This study has a 45-minute post-TNF treatment versus untreated comparison. Two DEG lists were generated: one conventional comparison looking at exonic RNA and another comparing intronic RNA (which can be used as a proxy for actively regulated genes (51)).

We applied DEET’s enrichment tool function to both the intronic- and exonic-calculated, TNFa-induced (upregulated) DEGs. We found that both intronic- and exonic-derived DEGs from Alizada *et al*., 2021 (51) retrieve comparisons related to TNFa treatment and bacterial infection (Figure 2A, Supplementary Figure S7). For example, the top 15 most enriched studies from each list include studies measuring gene expression after <1h of TNFa treatment (TNFa treatment to breast cancer cells for 40 minutes (54) and TNFa treatment to neutrophils for one hour (55)).

**Figure 2.**
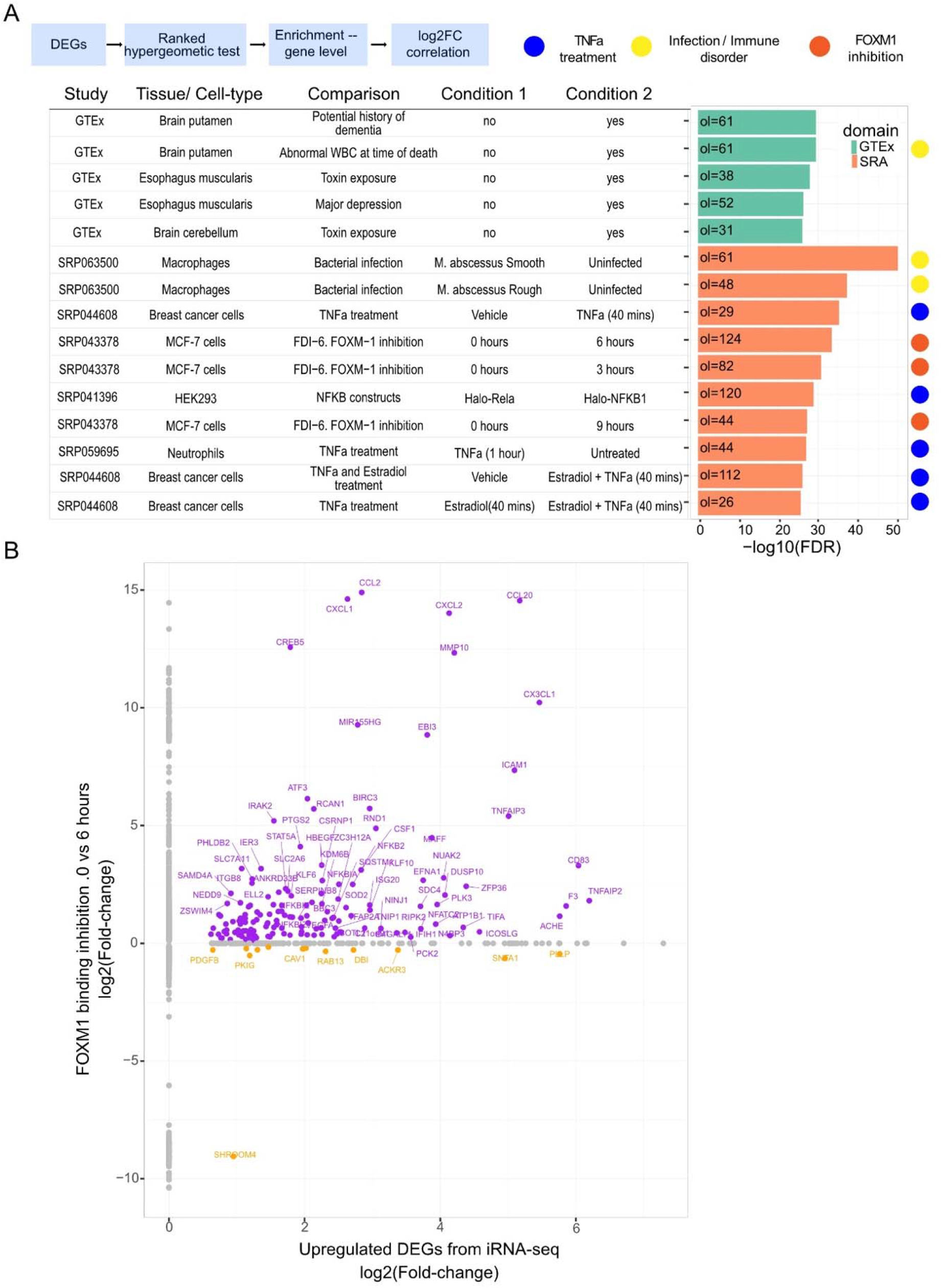
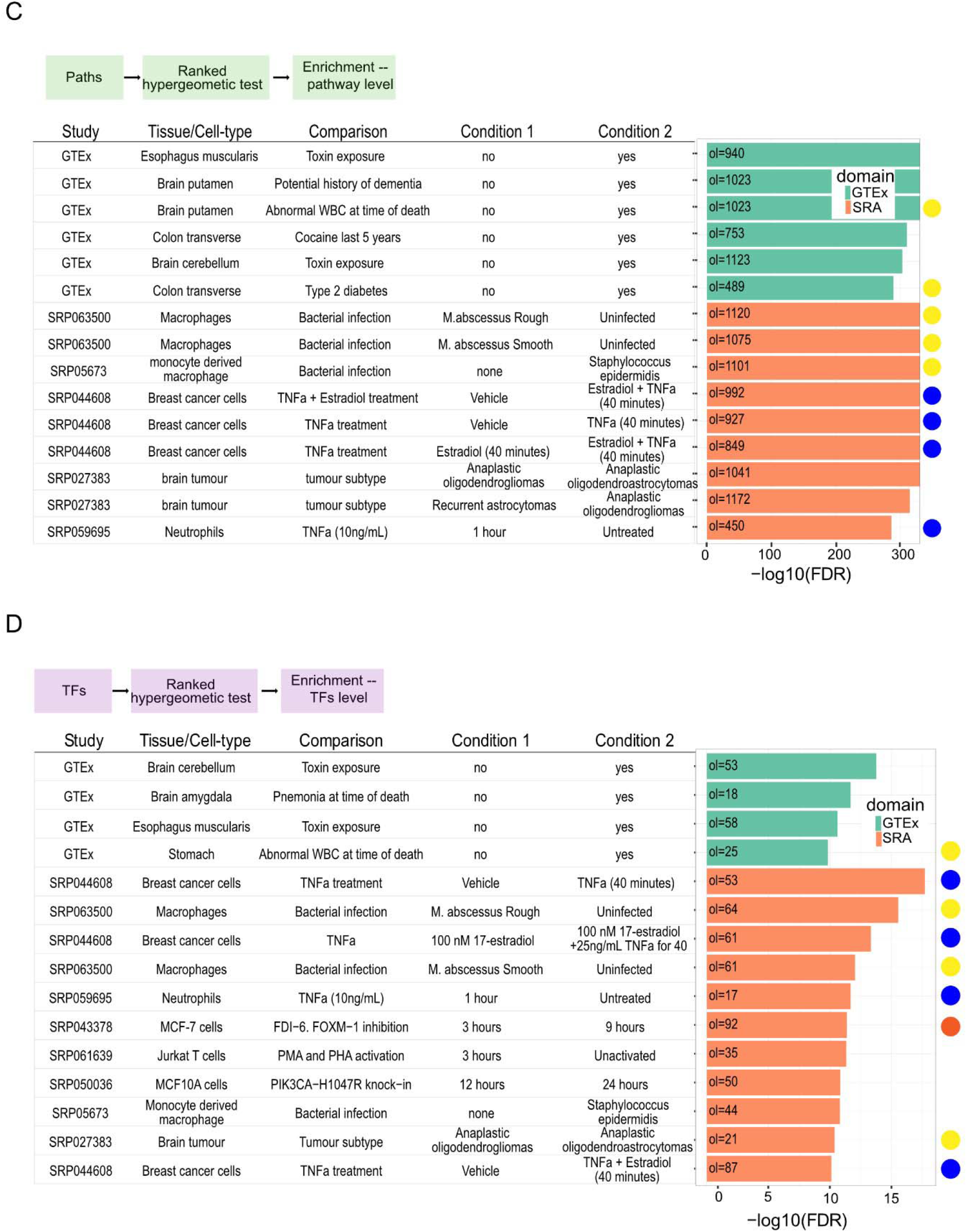
Summary of the DEET’s function applied to upregulated DEGs after TNFa treatment in Human Aortic endothelial cells (HAoECs) for 45 minutes from Alizada et al., 2021. A) Barplot of the top 15 most enriched pairwise comparisons based on overlapping DEGs from intronic RNA-seq. Rows are different comparisons within DEET, and the barplot is the - log10(FDR-adjusted p-value) of gene set enrichment computed by ActivePathways. B) Scatterplot of the log2(Fold-changes) of the upregulated DEGs in Alizada et al., 2021 from intronic RNA-seq (x-axis) vs. the DEGs in SRP043379 between 0 (naive) and 6 hours of FOXM1 inhibition (y-axis). Points are individual genes. Grey points are only DE in one study, purple points are DE in the same direction between studies, and orange points are DE in the opposite direction. C) Barplot of the top 10 most enriched pairwise comparisons based on overlapping biological pathways from intronic RNA-seq. Rows are different comparisons within DEET, and the barplot is the -log10(FDR-adjusted p-value) of path-set enrichment. D) Barplot of the top 10 most enriched pairwise comparisons based on overlapping TFs from intronic RNA-seq. Rows are different comparisons within DEET, and the barplot is the -log10(FDR-adjusted p-value) of the TF-set. For A, C, and D, comparisons annotated with a blue symbol are treatments of TNFa in different cell-lines. Comparisons annotated with a yellow symbol originate from infection and immune disorders studies. Comparisons annotated with an orange symbol originate from SRP043378, Gormally *et al*., 2014, which investigates differences in gene expression in MCF-7 after FOXM1 inhibition for 0 (naive) 3, 6, and 9 hours.

One important motivation for using DEET is to facilitate identifying new connections between one’s gene list and other studies that do not share a common experimental design. For example, the above DEET analysis of TNFa-treated endothelial cells returned a methods-based study looking at the effect of overexpressing NF-κB subunits RELA and NFKB1 in HEK293 cells (56) and another study of macrophages infected with *Mycobacterium abscesses* (57). We also retrieved a study that, at first glance, did not contain an obvious connection to proinflammatory gene responses but rather investigated differences in gene expression after FOXM1 inhibition in MCF7 breast cancer cells for 0 (naive) vs. 6 hours (39) (Figure 2A). We found a significant overlap of DEGs whose fold-changes were correlated (153 genes, R^2^ = 0.318, FDR-adjusted p-value = 1.849 × 10^−4^) (Figure 2B). While FOXM1 is often studied as a transcription factors that plays a role in proliferation and differentiation (39), previous studies link FOXM1 to TNF signaling through extensive chromatin co-localization of FOXM1 and NF-κB (58). These 153 overlapping genes significantly enrich the “TNFa signaling via NFkB” hallmark gene set (54 genes, FDR-adjusted p-value = 2.175 × 10^−66^). Lastly, DEET is also designed to identify significantly associated comparisons based on overlapping GO and TF-target terms obtained from user-submitted DEG lists. Using the above NF-κB DEG list and associated GO and TF-target terms, we identified additional DEG comparisons within the DEET dataset driven by GO terms “TNFa signaling via NF-κB”, “Response to lipopolysaccharide”, and “response to molecule of bacterial origin” (Figure 2C, D).

To further demonstrate the potential use of DEET and to provide an example where DEET was able to reveal novel biological insights that might be missed by transitional pathway enrichment analysis, we queried the list of genes downregulated after TNFa treatment. Such downregulated genes are known to have a weaker signal than upregulated genes and are often related to genes involved in cell-type-specific processes (59). We identified seven enriched comparisons using downregulated genes identified by integrating exonic and intronic DEGs (37). Interestingly, one comparison investigated breast cancer cells with both estradiol and TNFa treatment for 40 minutes (54), and another which investigated “11-18” lung adenocarcinoma cell line after pharmacological activation and inactivation of NF-κB (60) (Supplementary Figure S8). In contrast, traditional Gene Ontology enrichment (61) only identified pathways related to cell-lineage specificity (Supplementary Figure S8). We then investigated whether the seven overlapping genes between Alizada *et al*., 2021’s (51) downregulated genes and SRP044608 (estradiol + TNFa treatment) (54) have been previously linked to TNFa in the literature. Two overlapping genes, *TXNIP* (62) and *SMAD7* (63) are negatively correlated with TNFa treatment, and the other genes expressed based on TNFa varied based on the biological context (64–68).

### DEET identifies individual gene-gene associations across datasets

Lastly, DEGs that show correlated expression changes across different conditions are more likely to be part of the same biological pathway and undergo shared gene regulation (3, 69). We can leverage the associations of fold-changes between genes across all the comparisons in the DEET database to identify genes that may be under the same regulation. Specifically, the DEET_feature_extract() function detects genes associated with an input variable that can be assigned to every comparison (e.g., a gene of interest, whether the comparison investigated cancer, etc.) using an elastic net regression (45) in conjunction with correlation analysis to determine what genes are associated with the input variable. To showcase this application of DEET, we looked for genes whose fold changes are correlated with that of the TNFa encoding gene *TNF*. The 14 genes retrieved by DEET were enriched for “TNFa signaling via NFkB” more than any other gene ontology (FDR-adjusted p-value = 4.09 × 10^−11^) (Supplementary Figure S9A) and included well-known TNFa signaling genes *NFKBIA* (rank 2) and *SEMA4A* (rank 6) and (Supplementary Figure 9B).

Interestingly, the top-ranked gene was *CCDC7* (Supplementary 9B), a gene that is not annotated as a hallmark of TNFa signaling. Supporting the relevance of this hit, *CCDC7* has been shown to simultaneously activate interleukin-6 and the vascular endothelial growth factor (70), which TNFa can also do (71–73). Notably, comparisons within the DEET database where both *CCDC7* and *TNF* are DE did not include studies investigating short-term TNFa treatment. Instead, they included studies involving tumour vs. non-tumour, bacterial infection, and Crohn’s disease. Together, this vignette demonstrates how DEET can be used to obtain meaningful information from DEG comparisons made from uniformly processed public RNA-seq data.

## DISCUSSION

The Differential Expression Enrichment Tool (DEET) allows users to compare their DE gene lists to a curated atlas of 3162 DEG comparisons originating from GTEx, (16), TCGA (14), and studies within SRA (74). We envision DEET to be used alongside established and emerging tools that leverage uniformly processed data to allow users to discover biological patterns within their RNA-seq data (e.g. (7, 26, 27)).

A major challenge for implementing a tool like DEET, which investigates differential gene expression results in public data (29), lies in the scalability and consistency of publicly available metadata. We were able to build the DEET database because the PhenoPredict (19) tool annotated necessary metadata across every sample within SRA. However, there was considerable manual curation and study filtering even with this consistent annotation. The first major way to improve these annotations are with the continued development and use of metadata prediction algorithms like PhenoPredict (19), automated algorithms of existing metadata within SRA (8) like in MetaSRA (23) and *ffq* (https://github.com/pachterlab/ffq). The second major way to improve these annotations will be through community- and consortium-driven manual annotation of metadata such as the Biostudies and GEOMetaCuration tools (75) and (76). In the context of differential analysis, allowing researchers to report which variables are the experimental, stratifying, blocking, and covariate variables will be invaluable for tools like DEET to encompass larger uniformly processed datasets such as those provided by RNASeq-er (76), *recount3 (10)*, ARCHS4 (12), and refine.bio (https://www.refine.bio/) which collectively contains more RNA-seq studies from human and non-human species (10, 12).

Including model organism studies into differential gene expression databases is of great value given the greater diversity and controlled nature of study designs (i.e., tissue types, experimental variables, genetic backgrounds) which are not possible for human studies. In addition, public RNA-seq from model organisms will contain many smaller-scale, hypothesis-driven experiments compared to TCGA, and GTEx. Future developments of DEET would extend its database to searchable, consistently analyzed, and curated differential expression analyses collected from multiple species in Expression Atlas (27). Lastly, extending DEET to be able to search differential comparisons derived from consistent experiments beyond RNA-seq would be a logical next step to harness ongoing efforts for systematic analysis of public data from different genomic techniques such as scRNA-seq (20, 77), accessible chromatin profiling (ATAC-seq/DNAse-seq) (78, 79), and protein-DNA interactions mapping (ChIP-seq and in the future CUT&RUN/TAG) (80–83). In summary, by allowing users to rapidly connect their gene lists to a curated set of uniformly processed differential gene expression analyses, tools like DEET will facilitate access to the treasure trove of public RNA-seq data.

## DATA AVAILABILITY

### Supplementary Data

Code and data to regenerate the figures within this dataset can be found at figshare (https://doi.org/10.6084/m9.figshare.20427774.v1). Code and data to rebuild the DEET database can be found at figshare, however dbGap protected data in these code are excluded (https://doi.org/10.6084/m9.figshare.20425464.v1). A stable dataset of the DEET database at the time of submission can be found on zenodo (https://zenodo.org/record/6954162#.Yuv3f3bMI2w). The developmental dataset of the DEET database can be found at (http://wilsonlab.org/public/DEET_data).

## Author’s Contributions

D.S. conceived the method described in this manuscript. D.S. created the R code and the R package with support from J.A., K.E., H.H. Troubleshooting and methodological utility and testing of DEET tool was completed by D.S. with support from J.A., and K.E. The manuscript was written by D.S. with support from all authors. A.G. and M.W supervised the work. We would like to thank Derek Beaton, Leonardo Collado-Torres and Jeff Leek, and Gesse Gillis and Maggie Crow for their helpful discussions, as well as Sarah Watt for designing the DEET logo.

