## Supplemental Figures for "Differential Expression Enrichment Tool (DEET): An interactive atlas of human differential gene expression"

A

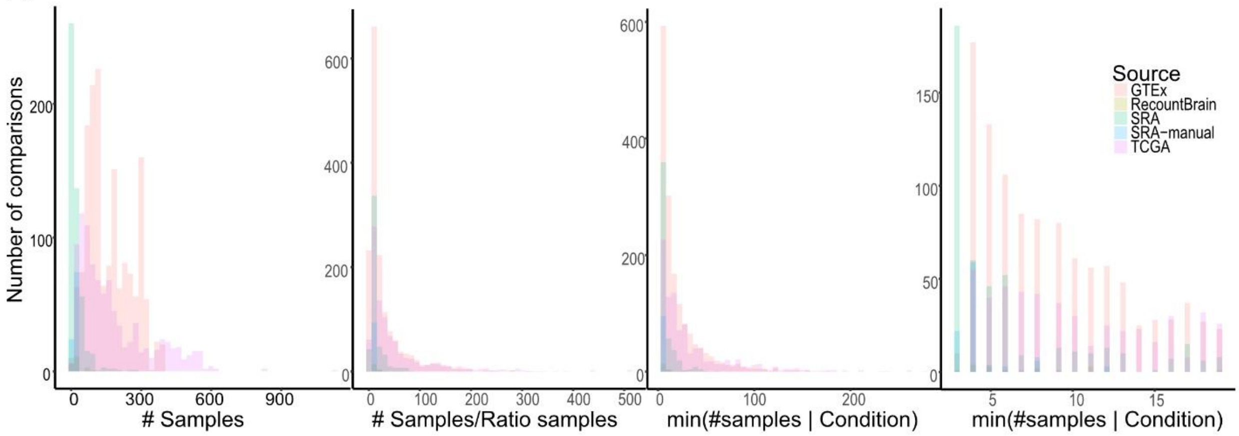


B

| Rho: log10(#DEGs) ~log10(#samples) | Rho: log10(#DEGs) ~ log10(#samples/RatioSamples) | Rho: log10(#DEGs) ~ log10(min(#Samples\|Condition) | Source |
|---|---|---|---|
| -0.435 | 0.289 | 0.238 | GTEx |
| -0.00689 | 0.145 | 0.468 | SRA |
| 0.504 | 0.399 | 0.450 | TCGA |

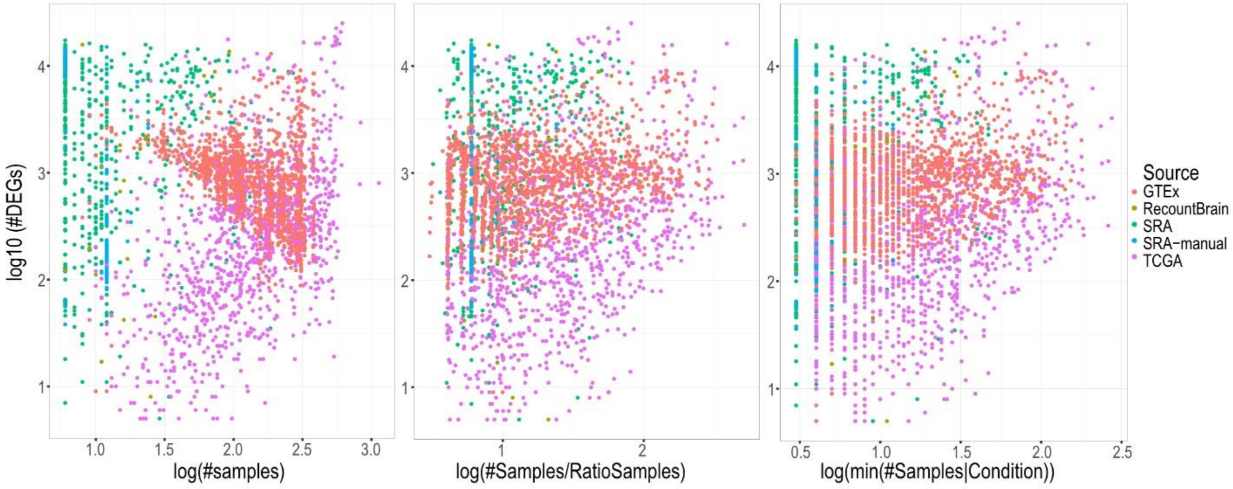

**Supplementary Figure S1.** Relationships between number of samples, number of comparisons, and number of differentially expressed genes (DEGs) across all comparisons within the DEET database. A) Histograms represent the distribution of sample numbers across the comparisons within the DEET database. Bar colour represents the comparison source (i.e., GTEx, SRA, SRA-manual, and TCGA). In all histograms, the Y-axis represents the number of comparisons. From left to right, the X-axes represent the number of samples in the comparison, the number of samples multiplied by the ratio of samples in condition 1 and condition 2, and the number of samples in the condition with fewer samples (i.e., minimum sample number). The final histogram increases resolution in minimum sample number by excluding comparisons with more than 20 samples in the condition with fewer samples. B) Scatterplots display the relationship between sample size and the number of DEGs. Each point is a comparison, colored by the comparison source (i.e., GTEx, SRA, SRA-manual, and TCGA). The Y-axis represents the log10(# DEGs) detected in each comparison. From left-to-right, the X-axes represent the number of samples in the comparison, the number of samples multiplied by the ratio of samples in condition 1 and condition 2, and the minimum sample number. The table above the scatterplots represents the correlation between # DEGs and sample size within each source.

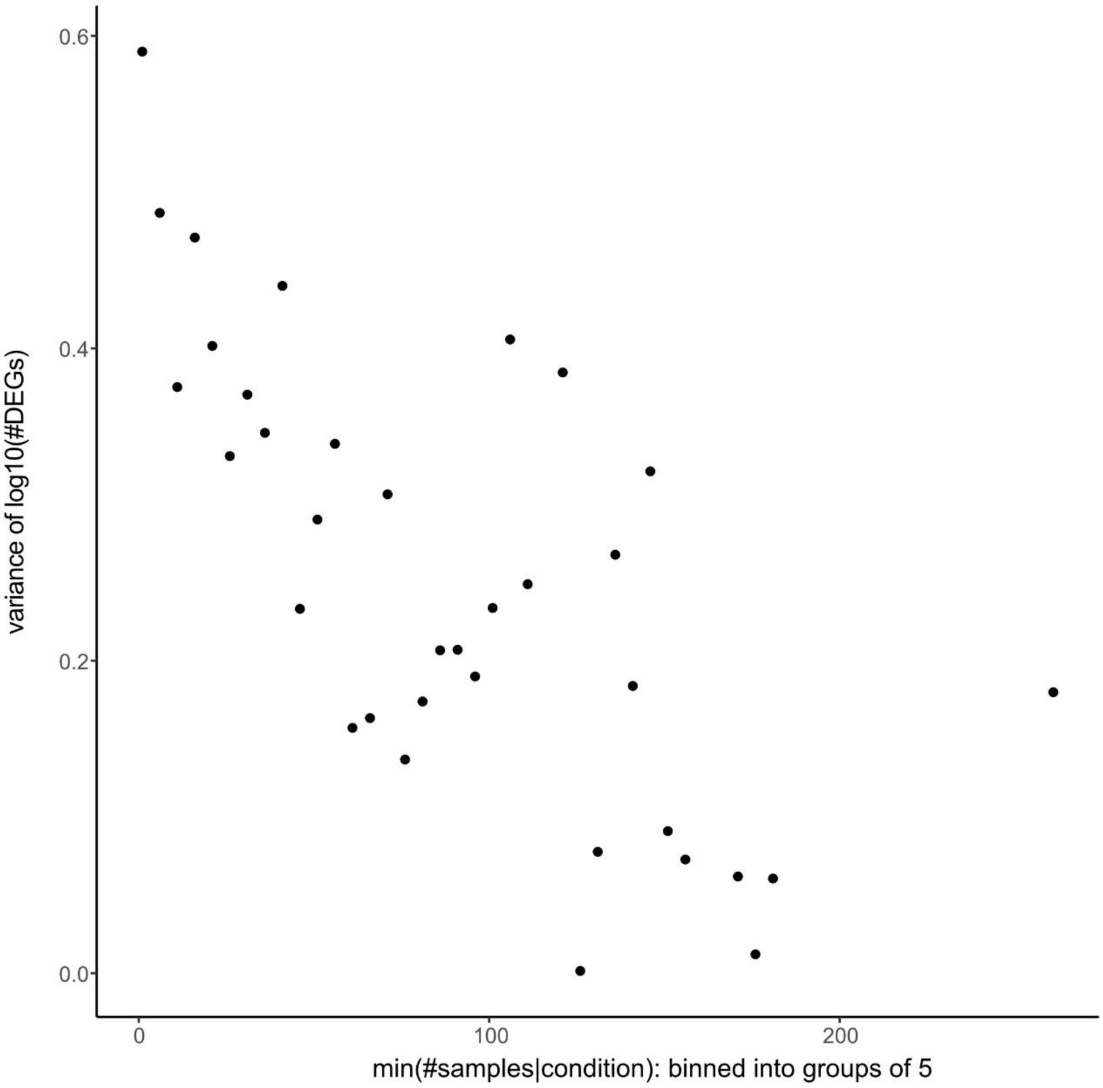


**Supplementary Figure S2.** Scatterplot displaying the relationship between the variance in differentially expressed genes (DEGs) and the number of samples in the condition with fewer samples (i.e., minimum sample number). We binned comparisons based on their minimum sample number into groups of 5 (e.g., conditions with a minimum sample number of 5-10 were binned) to be able to calculate the variance in # detected DEGs. The y-axis is the variance of the log10(# DEGs) within each bin. The x-axis is the lowest number in the minimum sample number bin (e.g., the 5-10 sample bin is plotted at 5 on the x-axis).

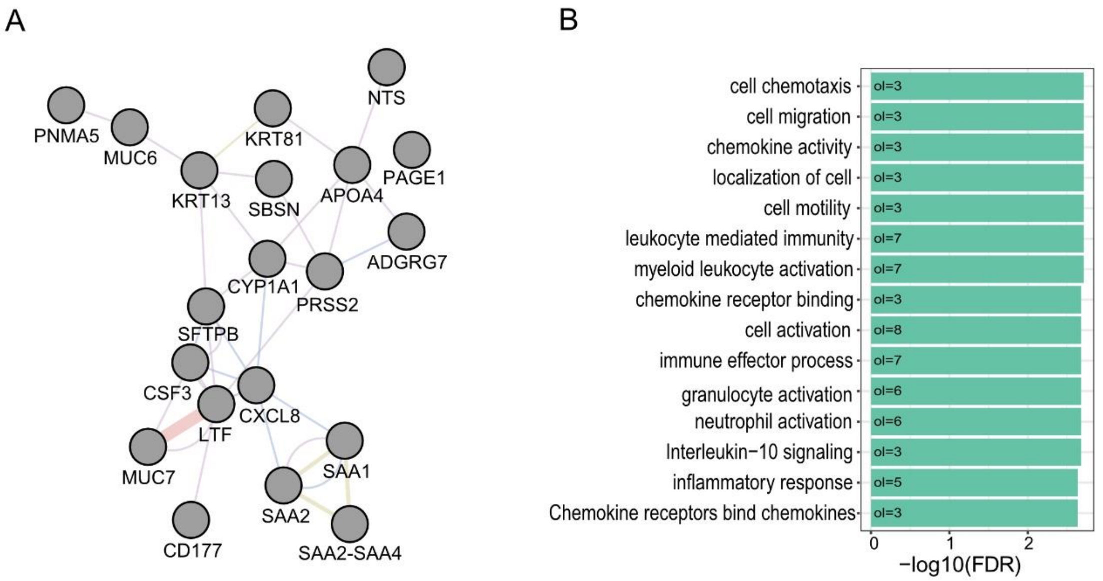


**Supplementary Figure S3.** Overlap between the top 1% of shared common DEGs in the DEET database and Crow *et al.,* 2019. A) Summary of the top 1% most common DEGs shared between the DEET database and Crow *et al.*, 2019. GeneMANIA plot of all common protein-coding genes (lncRNA TSIX and XIST are excluded in the plot because they are not in the GeneMANIA database). B) A barplot showing the gene-ontology enrichment of the genes in the top 1% most common DEGs in the DEET database and Crow *et al.*, 2019. The x-axis represents the -log10 False Discovery Rate. The number of genes overlapping "ol" with each gene ontology is shown within each bar.

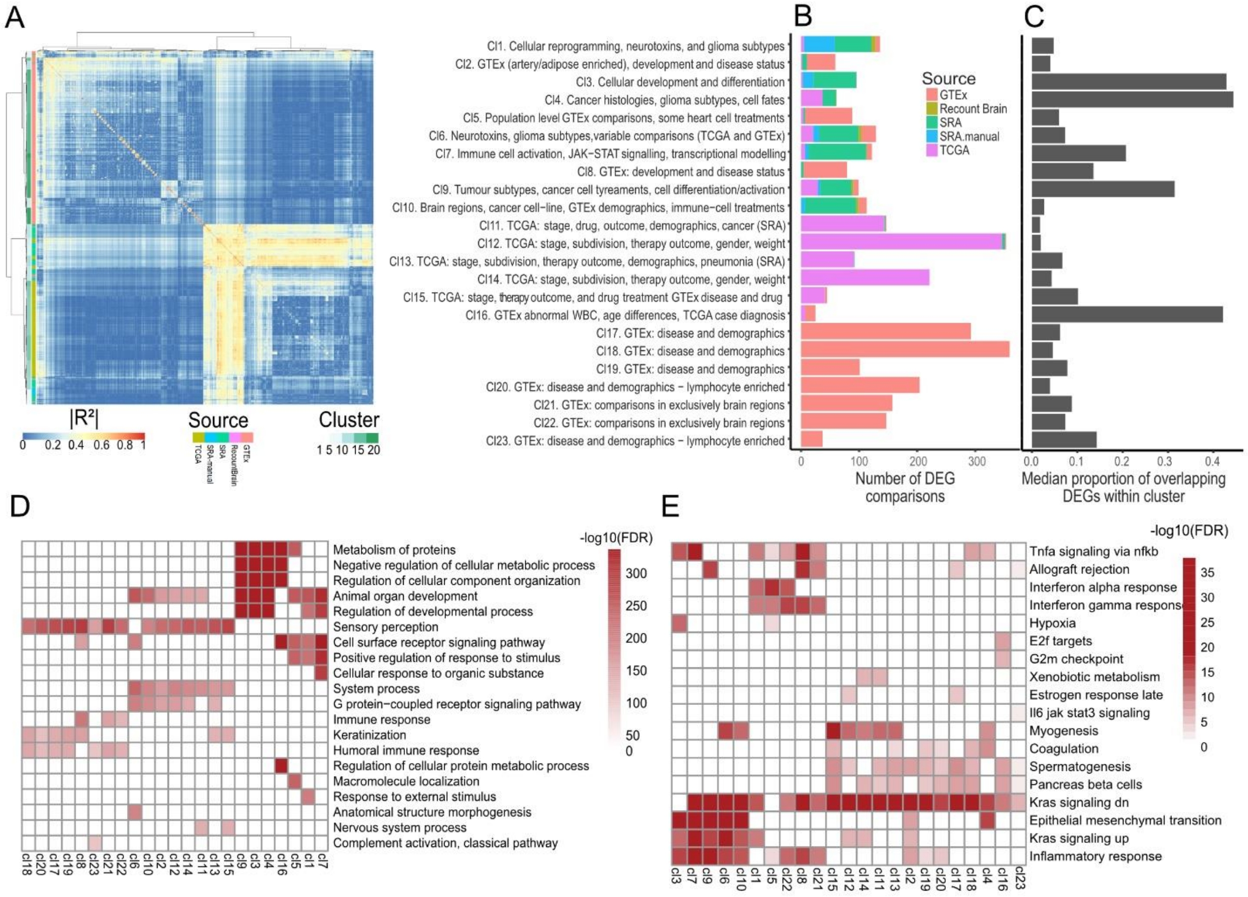


**Supplementary Figure S4.** Clustering and visualizing all pairwise comparisons within the Differential Expression Enrichment Tool (DEET). A) Pearson's correlation heatmap of all pairwise comparisons within the DEET database. For each pairwise comparison, the FDR-adjusted p-values of genes DE in at least one of the two comparisons were correlated. Rows and columns are comparisons and are clustered with Ward's D hierarchical clustering. Rows are annotated by cluster number and source (i.e., TCGA, SRA, and GTEx). B) Barplot of the number of comparisons within each cluster. Each cluster was labeled manually based on source, tissue, study design, and comparison type. Barplot height represents the number of comparisons in that cluster. Barplot colour represents the number of comparisons from each particular source. C) Barplot of the median proportion of DEGs shared within comparisons in each cluster. Clusters are labeled with the same summaries as in B. Bars represent the median proportion of overlapping genes that one comparison within a cluster will have with the other comparisons within that cluster. Gene Ontology (D) and Hallmark Gene Set (E) enrichment of each cluster of comparisons. In C) and D), only the top five pathways were plotted for each cluster. Rows are gene ontologies, and columns are clusters. The heatmap is populated with the -log10(FDR-adjusted p-value) of pathway enrichment. A gene ontology is only included for a cluster if it is in the top 10 most enriched pathways.

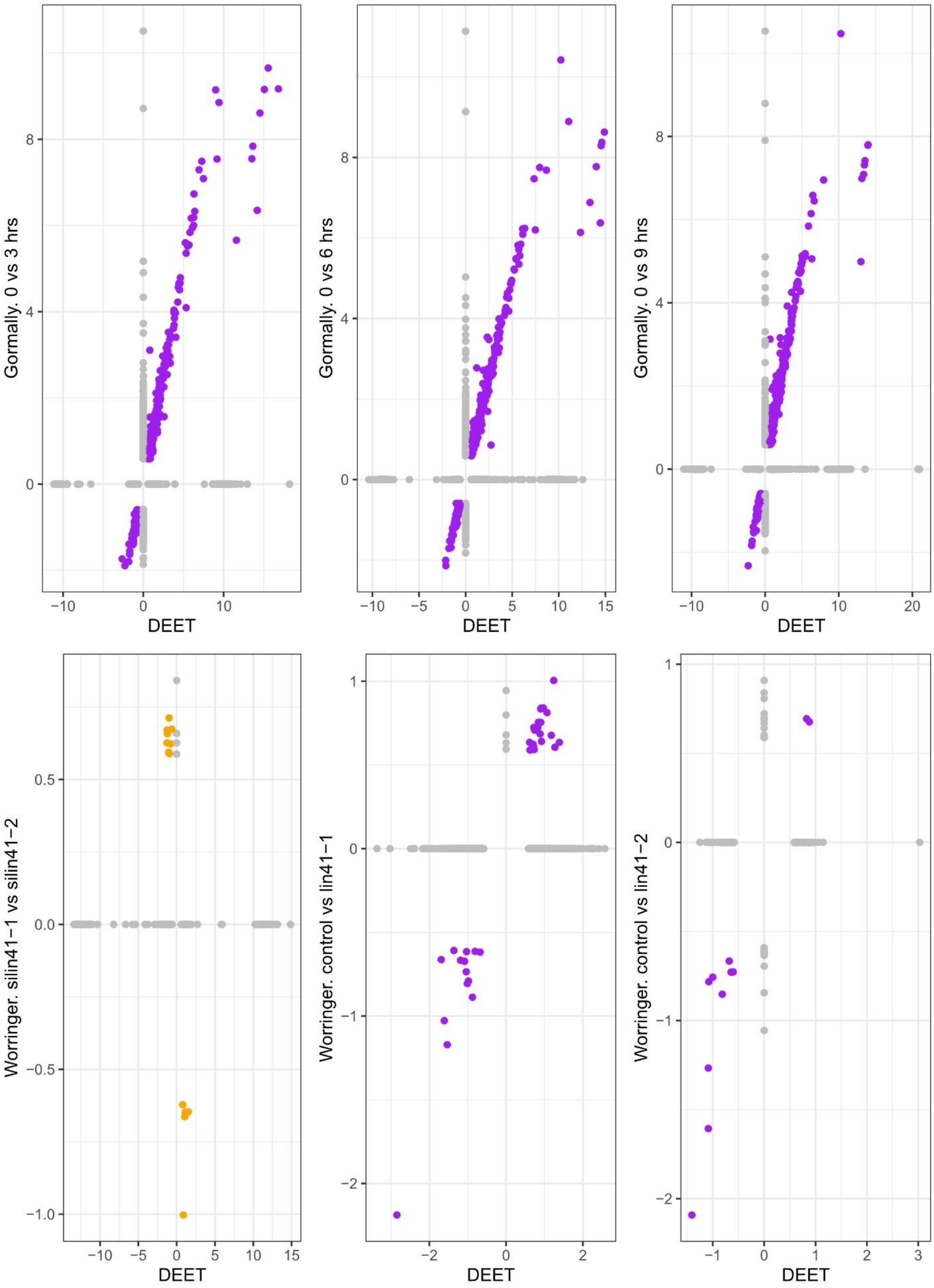
Gormally. 0 vs 3 hrs
Gormally. 0 vs 6 hrs
Gormally. 0 vs 9 hrs
Worringer. silin41−1 vs silin41−2
Worringer. control vs lin41−1
Worringer. control vs lin41−2
DEET

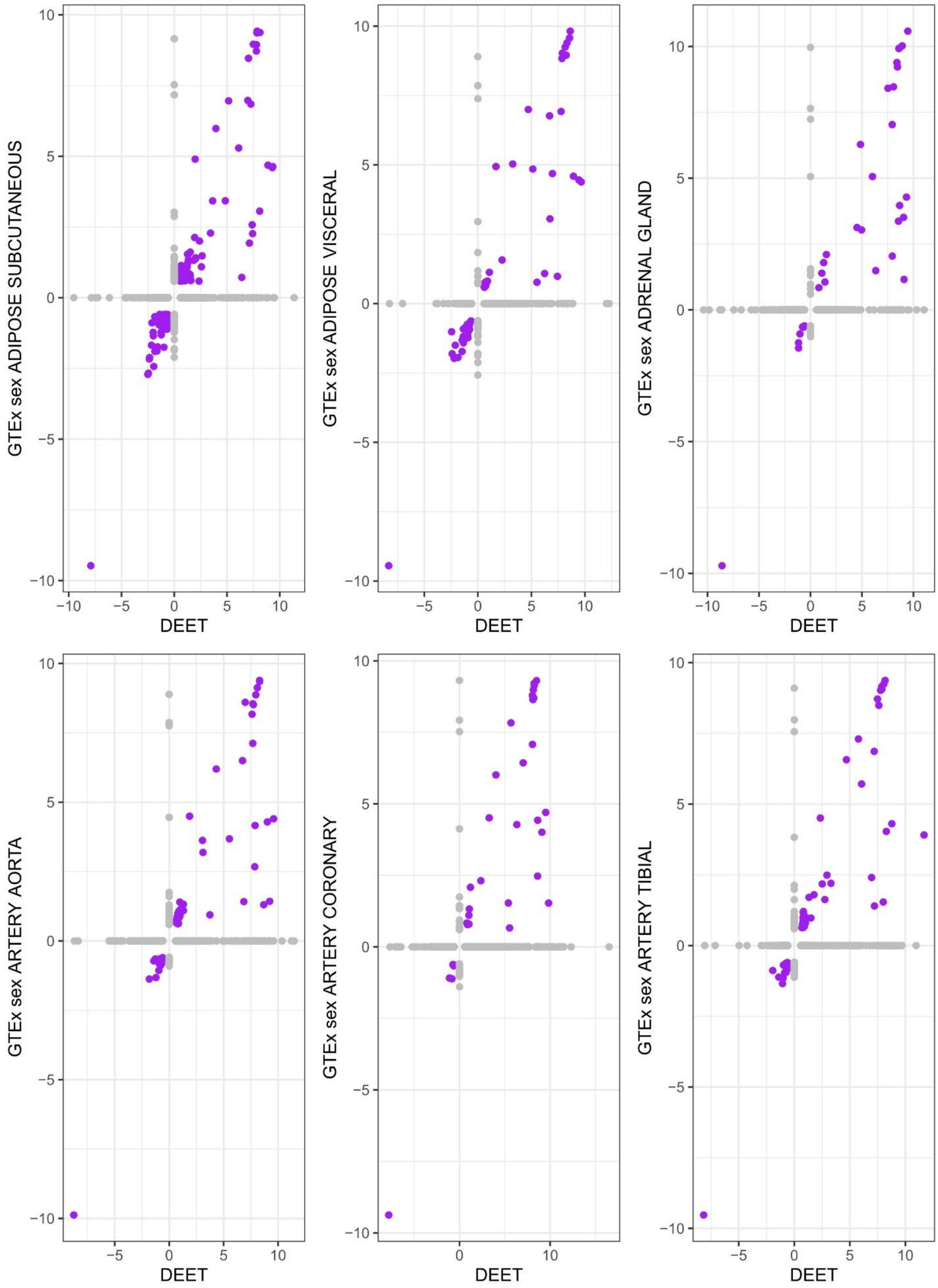
GTEx sex ADIPOSE SUBCUTANEOUS
DEET
GTEx sex ADIPOSE VISCERAL
DEET
GTEx sex ADRENAL GLAND
DEET
GTEx sex ARTERY AORTA
DEET
GTEx sex ARTERY CORONARY
DEET
GTEx sex ARTERY TIBIAL
DEET

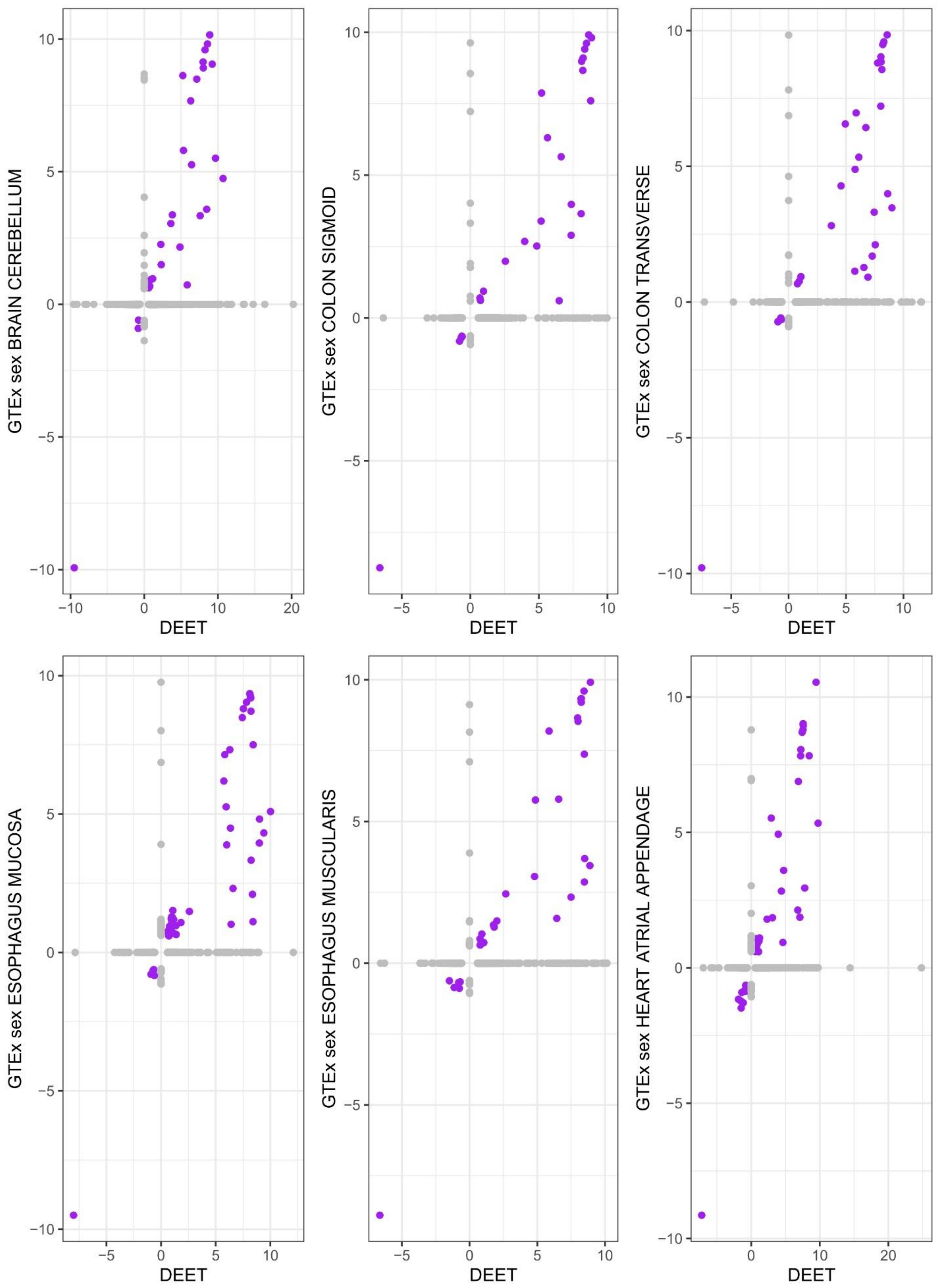

GTEx sex BRAIN CEREBELLUM
GTEx sex COLON SIGMOID
GTEx sex COLON TRANSVERSE
GTEx sex ESOPHAGUS MUCOSA
GTEx sex ESOPHAGUS MUSCULARIS
GTEx sex HEART ATRIAL APPENDAGE
DEET

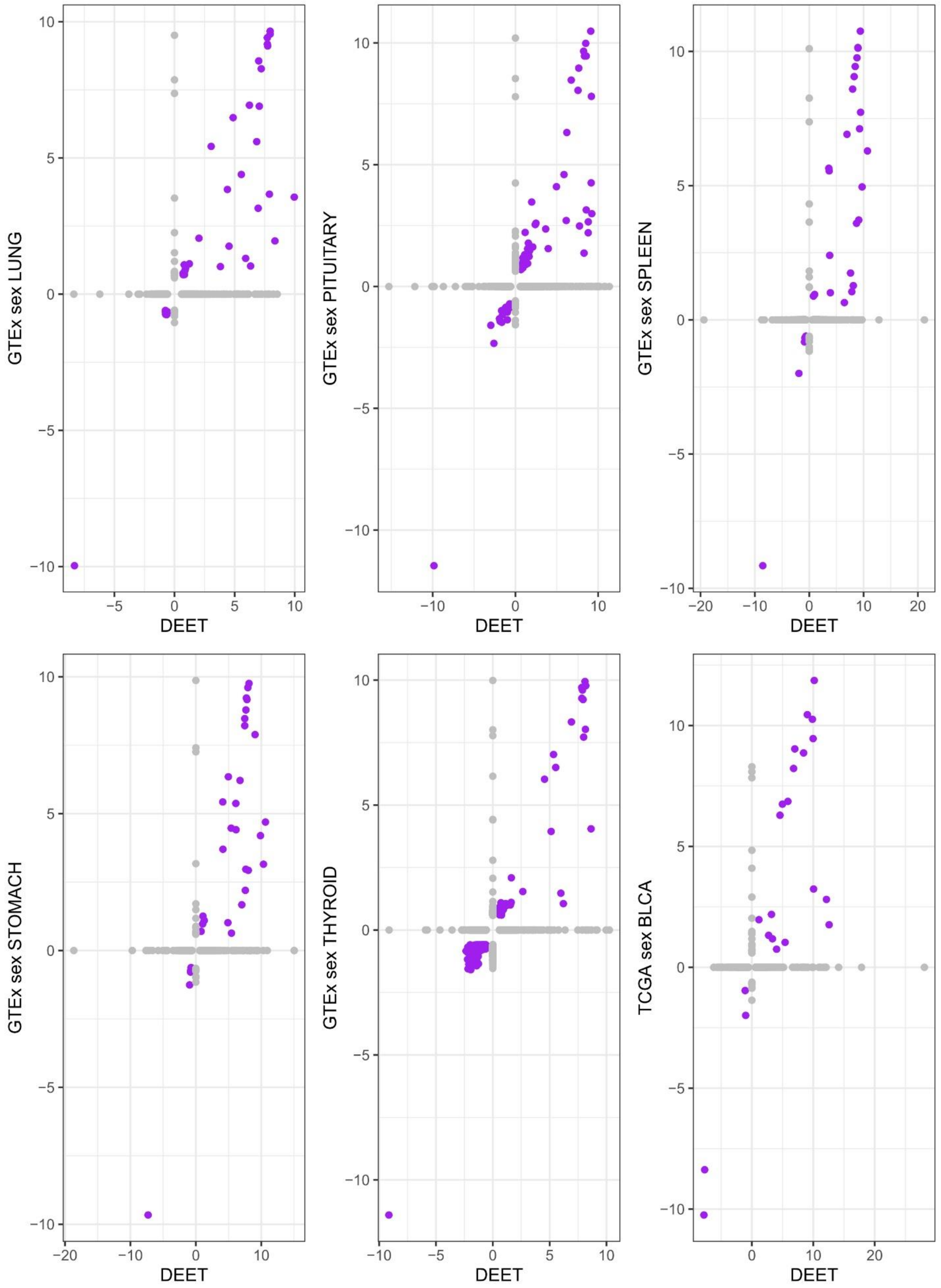
GTEx sex LUNG
GTEx sex PITUITARY
GTEx sex SPLEEN
GTEx sex STOMACH
GTEx sex THYROID
TCGA sex BLCA
DEET

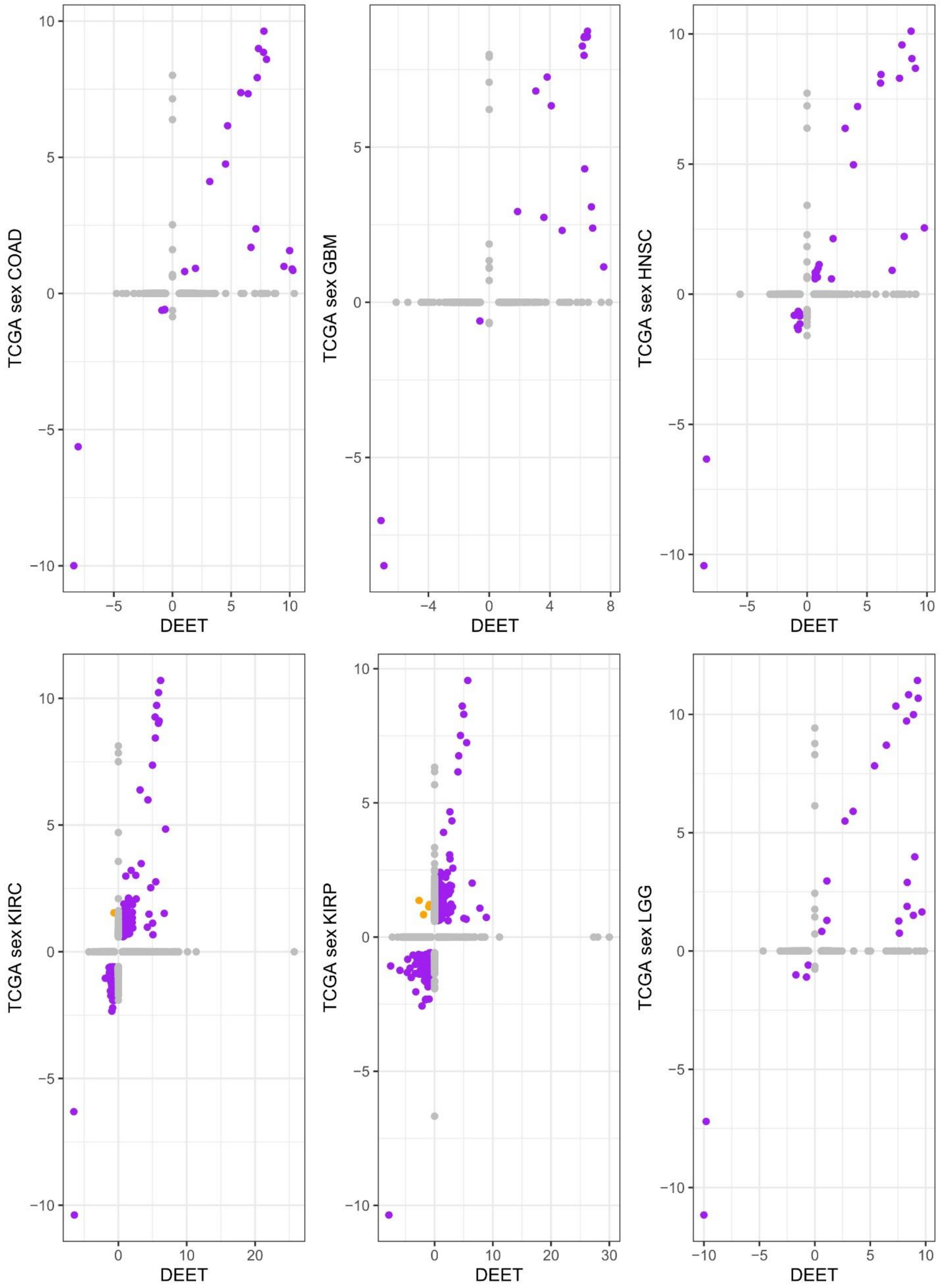
TCGA sex COAD
TCGA sex GBM
TCGA sex HNSC
TCGA sex KIRC
TCGA sex KIRP
TCGA sex LGG
DEET

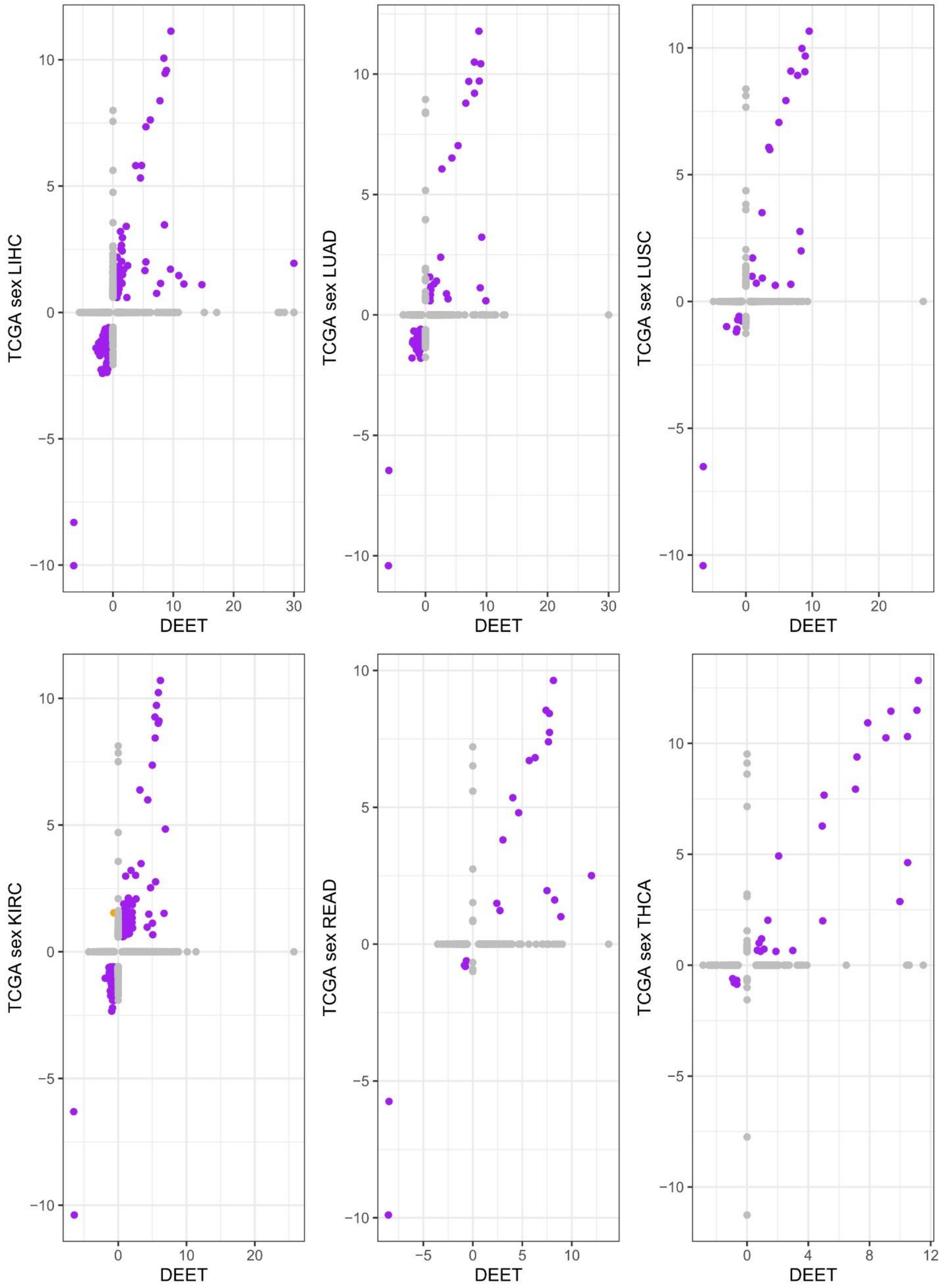
TCGA sex LIHC
DEET
TCGA sex LUAD
DEET
TCGA sex LUSC
DEET
TCGA sex KIRC
DEET
TCGA sex READ
DEET
TCGA sex THCA
DEET

**Supplementary Figure S5.** Scatterplots represent the log2(fold-changes) of differentially expressed genes (DEGs) within each benchmark comparison and their analogous comparison stored within the DEET database. Correlation coefficients for each pairwise comparison are stored in Table 1. The y-axis represents the log2(fold-changes) of genes from the benchmark comparison. The x-axis represents the log2(fold-changes) of the same gene from the analogous comparison stored within the DEET database. Each point is a DEG in the benchmark comparison or in the DEET database. Grey points are only DE in the benchmark study or the DEET database but not both. A gene that is only DE in the benchmark comparison is set to have an x-coordinate of 0, while a gene that is only DE in the analogous comparison in the DEET database is set to have a y-coordinate of 0. Purple points have the same fold-change within the benchmark comparison and the analogous comparison in the DEET database. Orange points contain the opposite fold-change between the benchmark comparison and the analogous comparison in the DEET database.

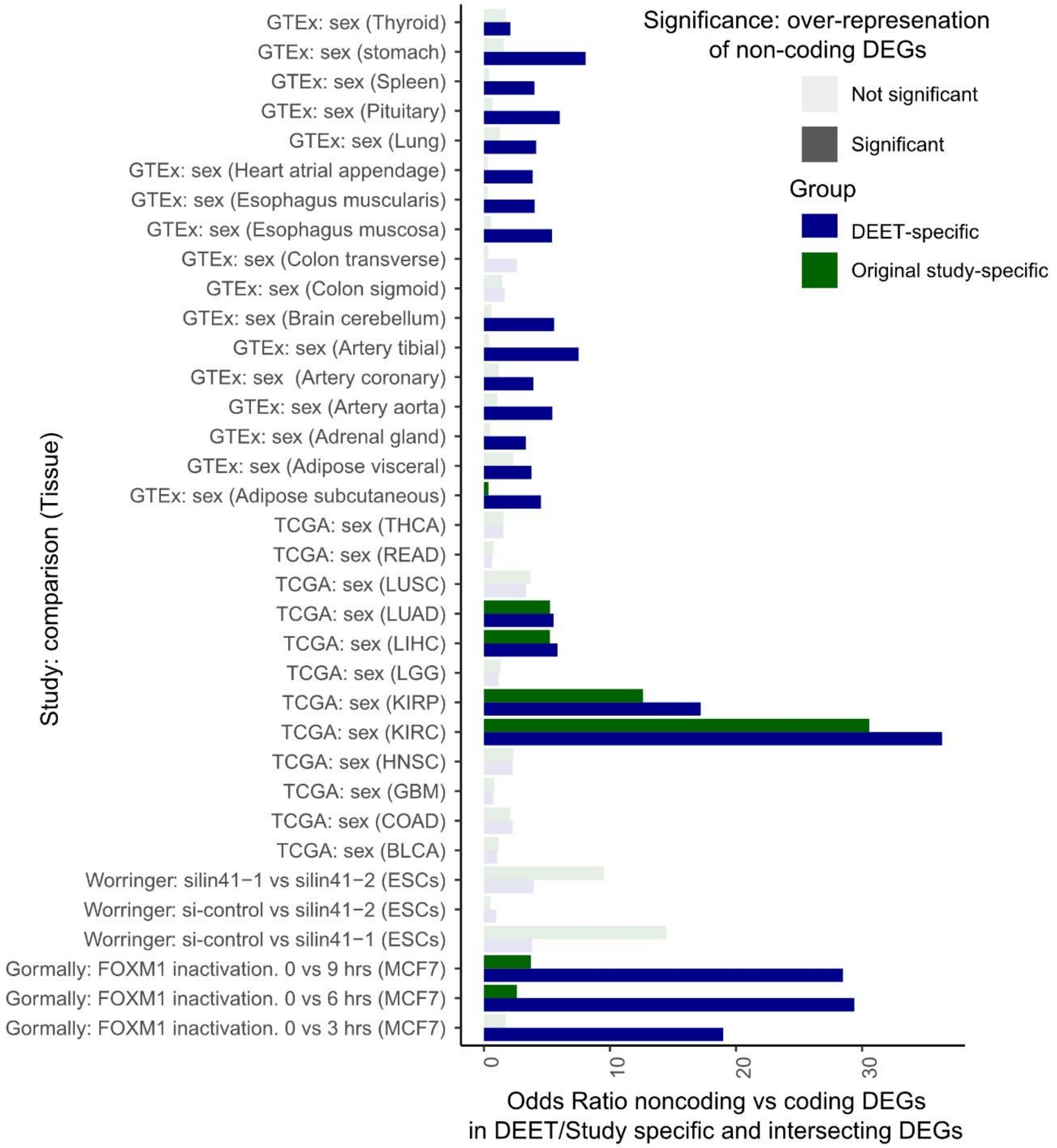


**Supplementary Figure S6.** Barplot representing the over-representation of non-coding DEET-specific and study-specific DEGs compared to DEGs shared between the DEET database and the original study (i.e., intersecting DEGS). Each row is a different benchmark comparison annotated by "data source. Comparison (tissue/cell)". The blue bar height represents the odds ratio of non-coding or coding genes in the DEET-specific vs. intersecting DEGs. The green bar height represents the odds ratio of non-coding or coding genes in the original study-specific vs.

intersecting DEGs. Opaque bars contain a significant over-representation of non-coding DEGs using a Fisher's exact-test (FDR-adjusted p-value $< 0.05$), and translucent bars are not significantly over-represented.

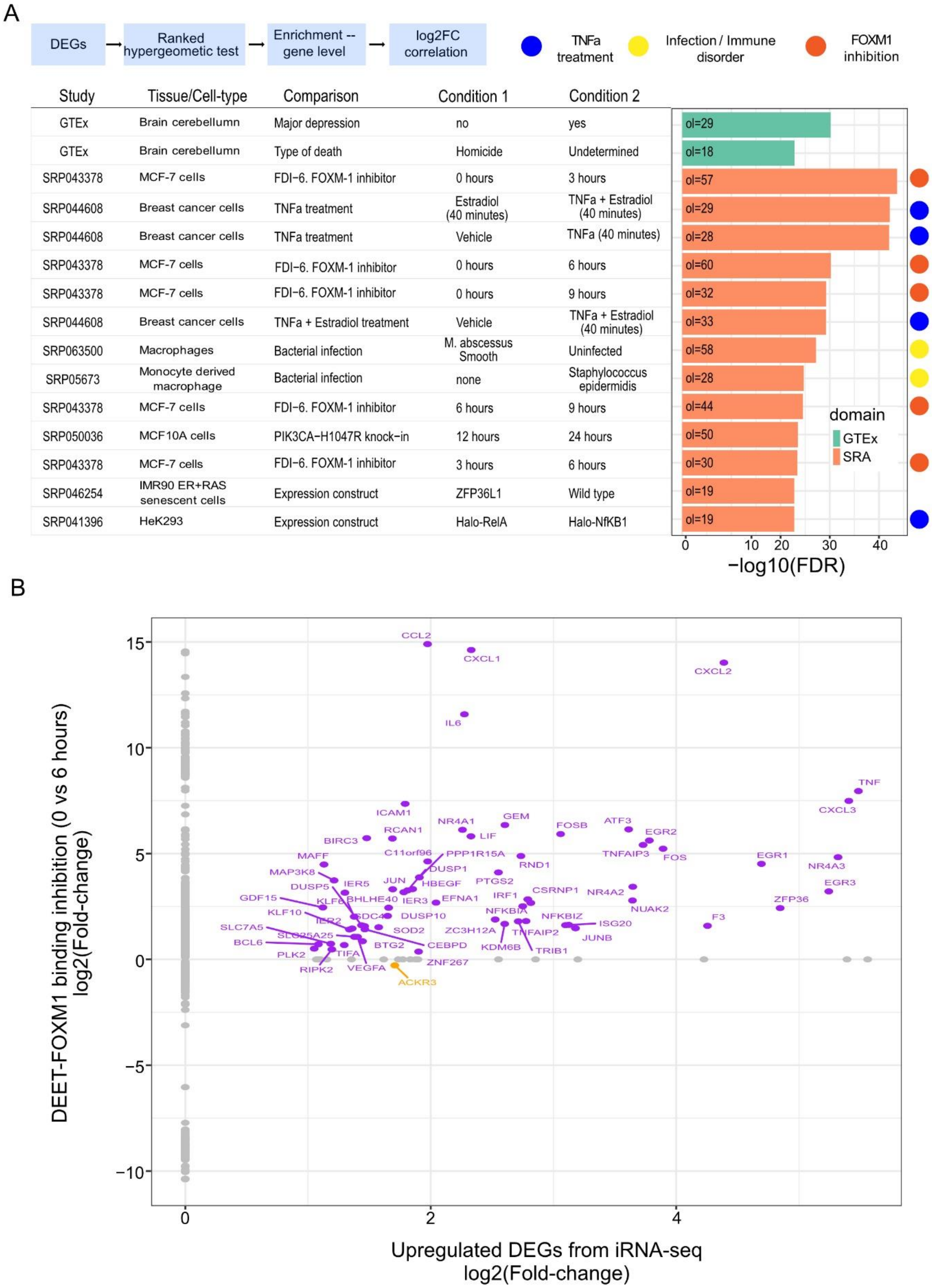
A
DEGs
Ranked hypergeometic test
Enrichment -- gene level
log2FC correlation
TNFa treatment
Infection / Immune disorder
FOXM1 inhibition
Study
Tissue/Cell-type
Comparison
Condition 1
Condition 2
GTEx Brain cerebellumn Major depression no yes ol=29
GTEx Brain cerebellumn Type of death Homicide Undetermined ol=18
SRP043378 MCF-7 cells FDI−6. FOXM-1 inhibitor 0 hours 3 hours ol=57
SRP044608 Breast cancer cells TNFa treatment Estradiol (40 minutes) TNFa + Estradiol (40 minutes) ol=29
SRP044608 Breast cancer cells TNFa treatment Vehicle TNFa (40 minutes) ol=28
SRP043378 MCF-7 cells FDI−6. FOXM-1 inhibitor 0 hours 6 hours ol=60
SRP043378 MCF-7 cells FDI−6. FOXM-1 inhibitor 0 hours 9 hours ol=32
SRP044608 Breast cancer cells TNFa + Estradiol treatment Vehicle TNFa + Estradiol (40 minutes) ol=33
SRP063500 Macrophages Bacterial infection M. abscessus Smooth Uninfected ol=58
SRP05673 Monocyte derived macrophage Bacterial infection none Staphylococcus epidermidis ol=28
SRP043378 MCF-7 cells FDI−6. FOXM-1 inhibitor 6 hours 9 hours ol=44
SRP050036 MCF10A cells PIK3CA−H1047R knock−in 12 hours 24 hours ol=50
SRP043378 MCF-7 cells FDI−6. FOXM-1 inhibitor 3 hours 6 hours ol=30
SRP046254 IMR90 ER+RAS senescent cells Expression construct ZFP36L1 Wild type ol=19
SRP041396 HeK293 Expression construct Halo-RelA Halo-NfKB1 ol=19
domain
GTEx
SRA
0 10 20 30 40
−log10(FDR)
B
DEET-FOXM1 binding inhibition (0 vs 6 hours) log2(Fold-change)
Upregulated DEGs from iRNA-seq log2(Fold-change)
15
10
5
0
−5
−10
0
2
4
CCL2
CXCL1
CXCL2
IL6
TNF
CXCL3
ICAM1
NR4A1
GEM
FOSB
ATF3
EGR2
RCAN1
LIF
BIRC3
C11orf96
PPP1R15A
TNFAIP3
FOS
EGR1
NR4A3
MAFF
DUSP1
RND1
MAP3K8
PTGS2
EGR3
DUSP5
IER5
JUN
HBEGF
CSRNP1
NR4A2
GDF15
KLF6
BHLHE40
IER3
EFNA1
IRF1
ZFP36
KLF10
SDC4
DUSP10
NFKBIA
NFKBIZ
NUAK2
F3
IER2
ISG20
SLC7A5
SOD2
ZC3H12A
TNFAIP2
SLC25A25
JUNB
BCL6
BTG2
CEBPD
KDM6B
TRIB1
PLK2
TIFA
ZNF267
RIPK2
VEGFA
ACKR3

**Supplementary Figure S7.** Summary of the DEET_enrich() function applied to upregulated DEGs after TNFa treatment in Human Aortic Endothelial Cells (HAoECs) for 45 minutes from Alizada *et al.*, 2021. A) Barplot of the top 15 most enriched pairwise comparisons based on overlapping DEGs from exonic RNA-seq. Rows are different comparisons within DEET, and the barplot is the -log10(FDR-adjusted p-value) of gene set enrichment computed by ActivePathways. B) Scatterplot of the log2(Fold-changes) of the upregulated DEGs in Alizada *et al.*, 2021 from exonic RNA-seq (x-axis) vs. the DEGs in SRP043379 between 0 (naive) and 6 hours of FOXM1 inhibition (y-axis). Points are individual genes. Grey points are only DE in one study, purple points are DE in the same direction between studies, and orange points are DE in the opposite direction. For A), comparisons annotated with a blue symbol are treatments of TNFa in different cell-lines. Comparisons annotated with a yellow symbol originate from infection and immune disorders studies. Comparisons annotated with an orange symbol originate from SRP043378, Gormally *et al.*, 2014, which investigates differences in gene expression in MCF-7 after FOXM1 inhibition for 0 (naive) 3, 6, and 9 hours.

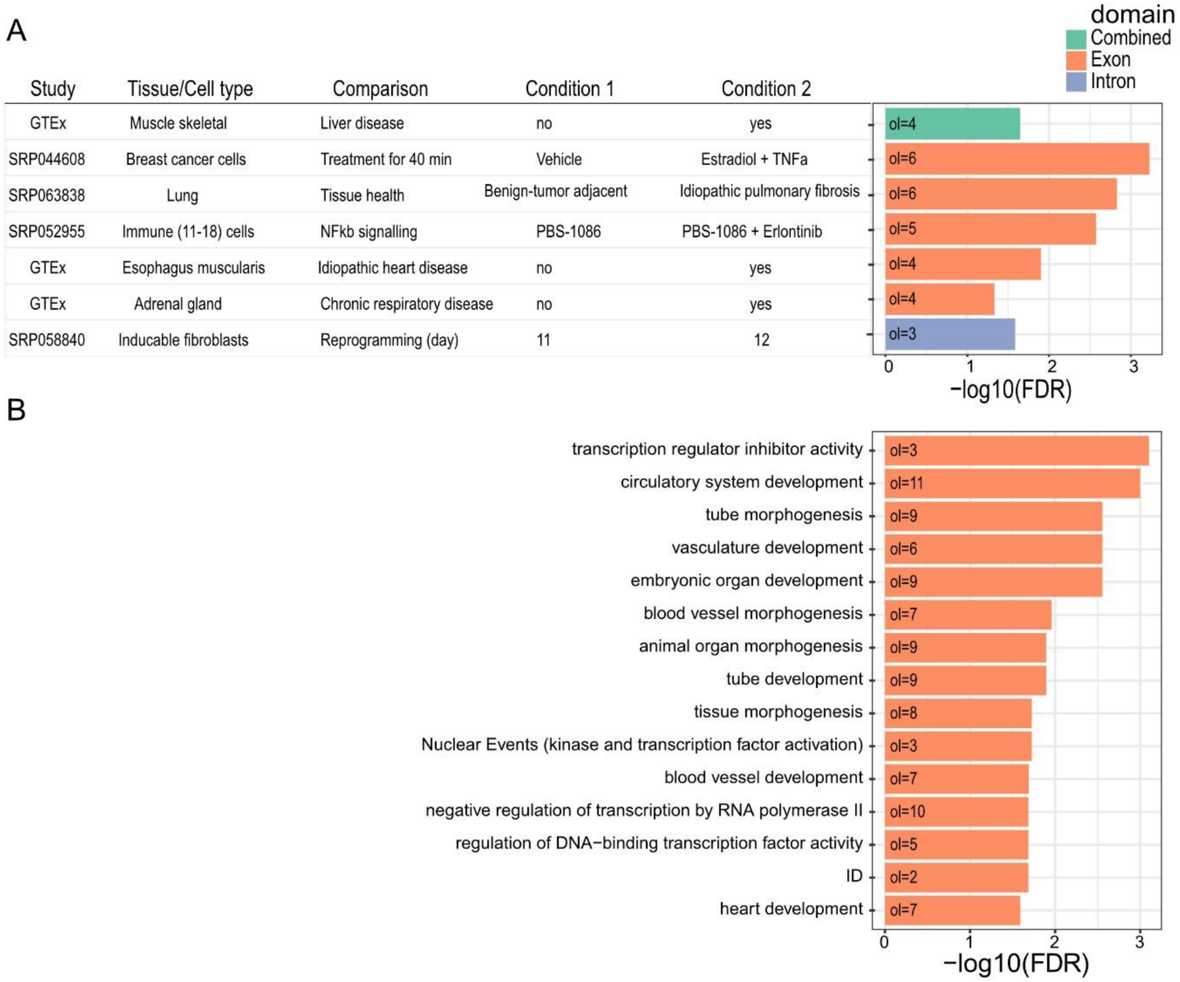


**Supplementary Figure S8.** Barplots representing the combined gene-set enrichment of downregulated DEGs from exonic RNA-seq and intronic RNA-seq using ActivePathways. FDR-adjusted p-values < 0.05 are considered significant. A) Barplot of enriched studies from the integrated downregulated DEGs using all DEG comparisons stored within DEET as the gene-set database. B) Barplot of enriched gene ontologies from the integrated downregulated DEGs using the traditional gene ontology database ("Human_GO_AllPathways_with_GO_iea_June_01_2021_symbo.gmt"). In both A) and B), Green bars demonstrate that enrichment was determined based on a combination of downregulated DEGs from exonic and intronic RNA-seq. Orange bars represent enrichment determined by downregulated DEGs from exonic RNA-seq. Blue bars represent enrichment determined by downregulated DEGs from intronic RNA-seq.

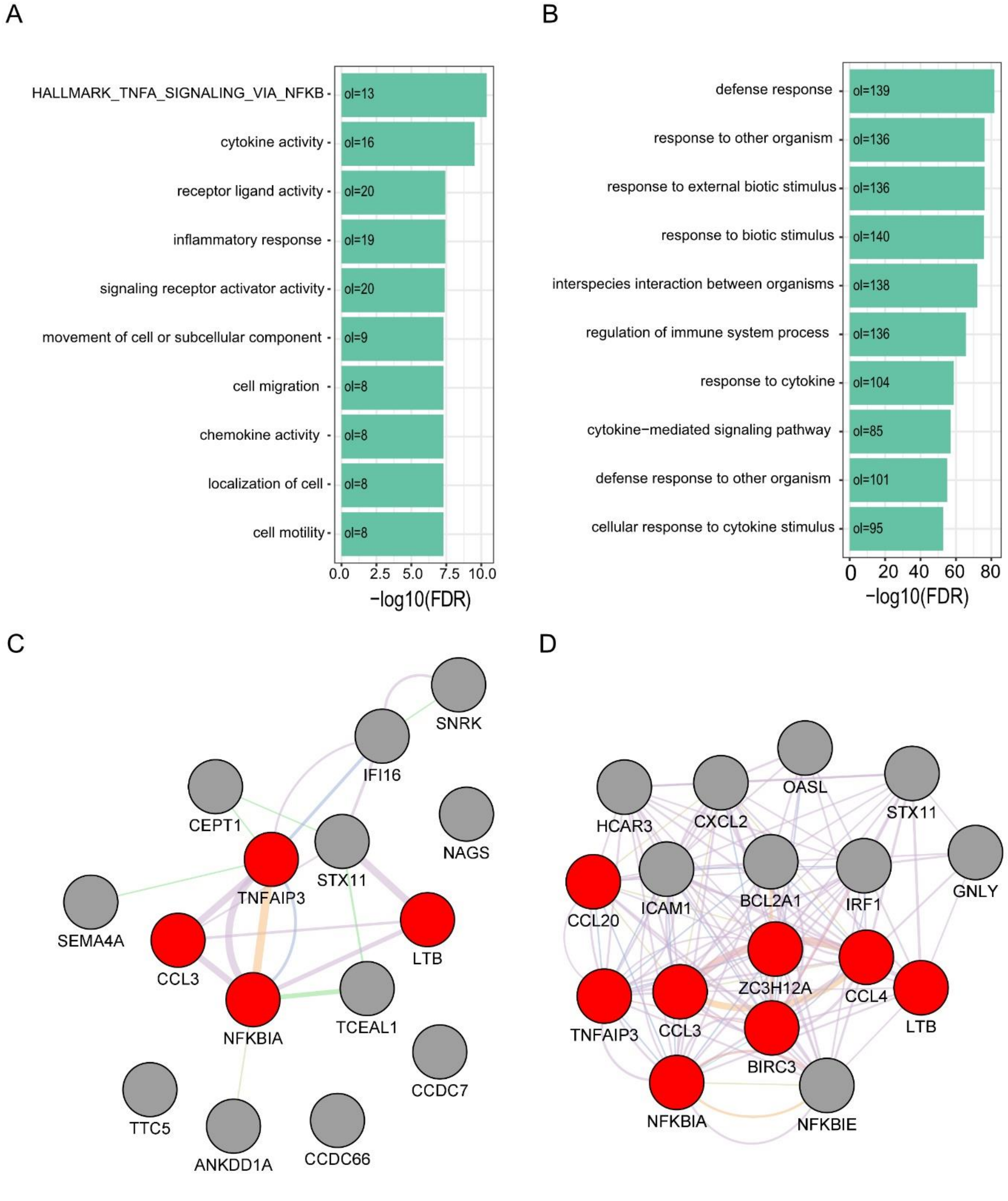


**Supplementary Figure S9**. Summary of genes with similar differential expression patterns (i.e., fold-changes) across DEET to the *TNF* gene. A) Barplot of gene ontology enrichment of genes whose differential expression is associated with *TNF* based on an elastic net regression. B) Barplot of gene ontology enrichment of genes whose differential expression is associated with *TNF* based on a Pearson's correlation. For A) and B), each row is a different enriched gene ontology, and the bar size is the -log10(FDR-adjusted p-value) of the hypergeometric gene set enrichment test. C) GeneMANIA barplot of the top 20 annotated genes whose differential expression pattern is most associated with *TNF* based on an elastic net regression. D) GeneMANIA barplot of the top 20 annotated genes whose differential expression pattern is most associated with *TNF* based on a Pearson's correlation analysis. For C) and D), genes are coloured in red if they are annotated to tumour necrosis factor response or tumour necrosis factor production pathways within the GeneMANIA database. Grey genes are not annotated to these pathways. Edge colours are the default datatype associations in GeneMANIA.
