## Supplemental File S2 for "Differential Expression Enrichment Tool (DEET): An interactive atlas of human differential gene expression"

### DEET Supplementary File S2

Dustin Sokolowski

20/07/2022

This workflow provides programatic examples of using the DEET CRAN package for the enrichment and visualization of the DEET database.

The enrichment and plotting components of DEET may also be used as a web-based tool here: https://wilsonlab-sickkids-uoft.shinyapps.io/DEET-shiny/ (https://wilsonlab-sickkids-uoft.shinyapps.io/DEET-shiny/)

#### Install DEET R package from CRAN

```
install.packages("DEET")
```

#### Require DEET R package from CRAN

```
library(DEET)
```

#### Read in the data required for all DEET functions.

All of these downloaded data are directly inputted into the functions within DEET. Accordingly, we reccomend saving these files once downloading them, that way you don't have to re-download them every time. Alternatively, these files can be downloaded directly from http://wilsonlab.org/public/DEET_data/ (http://wilsonlab.org/public/DEET_data/).

```
DEET_data <- DEET::DEET_data_download(x = "ALL")

DEET_dataset <- DEET_data$DEET_enrich
DEET_metadata <- DEET_data$metadata
DEET_features <- DEET_data$DEET_feature_extract
```

#### Run DEET enrich

With the DEET dataset downloaded, all that is required is to intput a list of genes (with or without associated fold-changes and p-values). There is also an optional background list of genes. In the context of RNA-seq, this is often all of the genes detected in an experiment. For this example, we will use the example gene-set built into the DEET R package with no background. Depending on how many comparisons are enriched, this typically takes 5-15 minutes on a laptop with 16Gb of RAM.

```
data(example_DEET_enrich_input)

# Running DEET with a list of genes including p-values and fold-changes
DEET_full_dataset <- DEET::DEET_enrich(DEG_list = example_DEET_enrich_input, DEET_dataset = D
EET_dataset,
                                      background = NULL)

# Running DEET with a generic list of genes including p-values and fold-changes

DEET_unordered_genes <- DEET::DEET_enrich(DEG_list = rownames(example_DEET_enrich_input), DEE
T_dataset = DEET_dataset,
                                      background = NULL, ordered = FALSE)
```

### Visualizing the outputs of DEET.

The files above have the same overall format. They are R lists with the following elements:

- AP_INPUT_BP_output: Output of traditional pathway enrichment of the gene list. Used to identify studies based on overlapping BPs.
- AP_INPUT_TF_output: Output of traditional TF gene-set enrichment of the gene list. Used to identify studies based on overlapping TFs.
- AP_DEET_DE_output: List containing the output of significantly enriched studies based on overlapping DEGs. It also contains the metadata for each of these studies.
- AP_DEET_BPs_output: List containing the output of significantly enriched studies based on overlapping BPs It also contains the metadata for each of these studies.
- AP_DEET_TF_output: List containing the output of significantly enriched studies based on overlapping TFss. It also contains the metadata for each of these studies.
- DE_correlations: List containing the correlation coefficients of enriched comparisons whose Fold-changes are ALSO correlated with the fold-changes of the inputted gene list. Note, this feature is not completed for the unordered gene list (since there are no co-efficients), so a message stating “No variance in coefs. Cannot proceed with correlation” is returned. This list also contains metadata for each of these enriched studies and dataframes providing the coefs of genes deemed signifcant in the input study and/or the enriched comparison.

```
# Print the top 6 enriched comparisons based on overlapping DEs.

head(DEET_full_dataset$AP_DEET_DE_output$results[order(DEET_full_dataset$AP_DEET_DE_output$re
sults$adjusted.p.val),])
```

```
##         term.id
## 1:  SRP043378_1
## 2: SRP044608_11
## 3:  SRP044608_2
## 4:  SRP043378_2
## 5:  SRP043378_3
## 6:  SRP063500_2
##
term.name
## 1:                          SRP043378: MC7 breast cancer. compound FDI-6 inhibiting bindin
g of FOXM-1. 0 hours (naive) vs 3 hours
## 2:             SRP044608: breast cancer cells. 100 nM 17-estradiol (E2) for 40min vs 100 nM
17-estradiol + 25 ng/mL TNFa for 40min
## 3:                                                   SRP044608: breast cancer cells. Vehic
le for 40min vs 25 ng/mL TNFa for 40min
## 4:                          SRP043378: MC7 breast cancer. compound FDI-6 inhibiting bindin
g of FOXM-1. 0 hours (naive) vs 6 hours
## 5:                          SRP043378: MC7 breast cancer. compound FDI-6 inhibiting bindin
g of FOXM-1. 0 hours (naive) vs 9 hours
## 6: SRP063500: Macrophages. bacterial infection - M. abscessus Smooth (MAB-S)-infected vs b
acterial infection - Uninfected control
##    adjusted.p.val term.size                                    overlap
## 1:   1.078194e-57      1688     TNFAIP3,ZFP36,CXCL3,SELE,JUNB,EGR1,...
## 2:   1.106540e-49       127   TNFAIP3,ZFP36,CXCL3,JUNB,EGR1,NFKBIZ,...
## 3:   4.209017e-48       125  TNFAIP3,CXCL3,SELE,JUNB,NFKBIZ,NFKBIA,...
## 4:   3.965164e-46      3748   TNFAIP3,ZFP36,CXCL3,JUNB,EGR1,NFKBIZ,...
## 5:   1.680232e-42      3756     TNFAIP3,ZFP36,CXCL3,SELE,JUNB,EGR1,...
## 6:   7.617565e-37       454 TNFAIP3,CXCL3,JUNB,NFKBIZ,CXCL2,NFKBIA,...
```

```
# Print the metadata of the top 6 enriched comparisons based on overlapping DEs.


head(DEET_full_dataset$AP_DEET_DE_output$metadata[order(DEET_full_dataset$AP_DEET_DE_output$r
esults$adjusted.p.val),])
```

```
##                    DEET.ID
## SRP043378_1   SRP043378_1
## SRP044608_11 SRP044608_11
## SRP044608_2   SRP044608_2
## SRP043378_2   SRP043378_2
## SRP043378_3   SRP043378_3
## SRP063500_2   SRP063500_2
##
DEET.Name
## SRP043378_1                            SRP043378: MC7 breast cancer. compound FDI-6 inhibit
ing binding of FOXM-1. 0 hours (naive) vs 3 hours
## SRP044608_11            SRP044608: breast cancer cells. 100 nM 17-estradiol (E2) for 40min
vs 100 nM 17-estradiol + 25 ng/mL TNFa for 40min
## SRP044608_2                                                 SRP044608: breast cancer ce
lls. Vehicle for 40min vs 25 ng/mL TNFa for 40min
## SRP043378_2                            SRP043378: MC7 breast cancer. compound FDI-6 inhibit
ing binding of FOXM-1. 0 hours (naive) vs 6 hours
## SRP043378_3                            SRP043378: MC7 breast cancer. compound FDI-6 inhibit
ing binding of FOXM-1. 0 hours (naive) vs 9 hours
## SRP063500_2  SRP063500: Macrophages. bacterial infection - M. abscessus Smooth (MAB-S)-inf
ected vs bacterial infection - Uninfected control
##
Description
## SRP043378_1                                  Suppression of the FOXM1 transcriptional progr
am via novel small molecule inhibition
## SRP044608_11                    TNFa Signaling Exposes Latent Estrogen Receptor Binding 
Sites in Breast Cancer Cells [GRO-seq]
## SRP044608_2                     TNFa Signaling Exposes Latent Estrogen Receptor Binding 
Sites in Breast Cancer Cells [GRO-seq]
## SRP043378_2                                  Suppression of the FOXM1 transcriptional progr
am via novel small molecule inhibition
## SRP043378_3                                  Suppression of the FOXM1 transcriptional progr
am via novel small molecule inhibition
## SRP063500_2  High-throughput RNA-sequencing of human macrophages infected with Mycobacteri
um abscessus Smooth and Rough variants
##              Source  Study.ID Samples
## SRP043378_1     SRA SRP043378       6
## SRP044608_11    SRA SRP044608       6
## SRP044608_2     SRA SRP044608       6
## SRP043378_2     SRA SRP043378       6
## SRP043378_3     SRA SRP043378       6
## SRP063500_2     SRA SRP063500      14
##                                                                   Samples.Up
## SRP043378_1                                                     time: 3 h:3
## SRP044608_11 treated with: 100 nM 17ß-estradiol + 25 ng/mL TNFa for 40min:3
## SRP044608_2                                 treated with: Vehicle for 40min:3
## SRP043378_2                                                     time: 6 h:3
## SRP043378_3                                                     time: 9 h:3
## SRP063500_2     bacterial infection: M. abscessus Smooth (MAB-S)-infected:6
##                                                      Samples.Down  Tissue numDEs
## SRP043378_1                                        time: 0 h:3 Bladder   1688
## SRP044608_11 treated with: 100 nM 17ß-estradiol (E2) for 40min:3  Breast    127
## SRP044608_2             treated with: 25 ng/mL TNFa for 40min:3  Breast    125
## SRP043378_2                                        time: 0 h:3 Bladder   3748
## SRP043378_3                                        time: 0 h:3 Bladder   3756
```

```
## SRP063500_2            bacterial infection: Uninfected control:8  Uterus    454
##              numDEup numDEdown Age.Mean.SD    Sexes
## SRP043378_1      981       707             female 6
## SRP044608_11     108        19             female 6
## SRP044608_2        7       118             female 6
## SRP043378_2     2155      1593             female 6
## SRP043378_3     2230      1526             female 6
## SRP063500_2       80       374               male 14
##
DEGs.Up
## SRP043378_1                LIF;RCAN1;EGR3;FOS;NR4A3;CXCL8;FOSB;TNFAIP3;ZFP36;PPP1R15A;TNF;
NFKBIA;RASD1;ATF3;NR4A1
## SRP044608_11              BIRC3;NFKBIA;EGR3;TNFAIP3;RELB;EGR1;CD83;MAFF;NFKB2;CXCL3;TNF;NF
KBIZ;CXCL8;NFKBID;CXCL1
## SRP044608_2                                                SNORD95;AVPR1A;C1orf21-DT;CHKB-D
T;TCHH;LYSMD1;LINC02188
## SRP043378_2       LIF;RCAN1;FOS;NR4A3;HMOX1;CYP1B1;CYP1A1;SLC7A11;TNFAIP3;ZFP36;PPP1R15A;T
NF;NFKBIA;HSPA1A;HSPA1B
## SRP043378_3  LIF;RCAN1;FOS;NR4A3;HMOX1;CYP1B1;CYP1A1;SLC7A11;TNFAIP3;PPP1R15A;HERPUD1;NFKB
IA;HSPA1A;HSPA1B;DNAJB1
## SRP063500_2        PGF;LYL1;HDDC3;P3H4;WDR88;DOK2;ADGRG6;AKNA;RIMBP3C;INKA1;TMEM129;SNAI3;
SH3TC1;DIPK1B;LRRC37A6P
##
DEGs.Down
## SRP043378_1                       KIF20A;PLK1;AURKA;SHH;PSRC1;CDC20;PRR11;CCNB1;TPX2;NEURL
1B;KPNA2;BUB1;TOP2A;CCNF;TROAP
## SRP044608_11          AMH;HMGN1P15;NBEAL2;RGL2;INPPL1;RHPN1;TCHH;FOXH1;KIFC2;TRABD;RELL2;S
GSM3;SLC25A29;ULK3;KIAA0895LP1
## SRP044608_2  TNFAIP3;CXCL3;BIRC3;MAP3K8;EFNA1;TNFRSF11B;NFKBIA;NFKB2;RIPK2;PHLDB2;NR4A1AS;
CACNG8;TICAM1;RAB3IP;THBS1-IT1
## SRP043378_2              KIF20A;PLK1;CDC20;SHH;AURKA;CACNG4;CCNB1;CCND1;DLGAP5;CXXC5;PSRC
1;ARL6IP1;KNSTRN;PRKCD;SLC9A3R1
## SRP043378_3      IGFBP5;SLC9A3R1;SPDEF;PREX1;KIF20A;CACNG4;TMEM164;ZNF467;SDC1;KANK2;ESRP
2;PSRC1;CXXC5;EPN3;NPHP3-ACAD11
## SRP063500_2             CXCL2;TNF;CCL4L2;CCL4;CXCL1;IER3-AS1;CD274;IL1A;SOCS3;IER3;CCL3-AS
1;SAT1;TNFAIP3;CXCL3;MIR3142HG
##
BP.sig
## SRP043378_1
SUPERPATHWAY OF CHOLESTEROL BIOSYNTHESIS%HUMANCYC%PWY66-5;ANDROGENRECEPTOR%IOB%ANDROGENRECEPT
OR;CCR1%IOB%CCR1;CD40%IOB%CD40;CXCR4%IOB%CXCR4
## SRP044608_11
CD40%IOB%CD40;CXCR4%IOB%CXCR4;EGFR1%IOB%EGFR1;EPO%IOB%EPO;GDNF%IOB%GDNF
## SRP044608_2
CD40%IOB%CD40;CXCR4%IOB%CXCR4;GDNF%IOB%GDNF;GM-CSF%IOB%GM-CSF;TNFALPHA%IOB%TNFALPHA
## SRP043378_2                       SUPERPATHWAY OF INOSITOL PHOSPHATE COMPOUNDS%HUMANCYC%PW
Y-6371;TRNA CHARGING%HUMANCYC%TRNA-CHARGING-PWY;3-PHOSPHOINOSITIDE BIOSYNTHESIS%HUMANCYC%PWY-
6352;SUPERPATHWAY OF CHOLESTEROL BIOSYNTHESIS%HUMANCYC%PWY66-5;ALPHA6BETA4INTEGRIN%IOB%ALPHA6
BETA4INTEGRIN
## SRP043378_3  SUPERPATHWAY OF INOSITOL PHOSPHATE COMPOUNDS%HUMANCYC%PWY-6371;PHOSPHATIDYLGL
YCEROL BIOSYNTHESIS II (NON-PLASTIDIC)%HUMANCYC%PWY4FS-8;TRNA CHARGING%HUMANCYC%TRNA-CHARGING
-PWY;CDP-DIACYLGLYCEROL BIOSYNTHESIS I%HUMANCYC%PWY-5667;3-PHOSPHOINOSITIDE BIOSYNTHESIS%HUMA
NCYC%PWY-6352
## SRP063500_2
CCR1%IOB%CCR1;CD40%IOB%CD40;CXCR4%IOB%CXCR4;EGFR1%IOB%EGFR1;FLK2 FLT3%IOB%FLK2 FLT3
##
```

```
TF.sig
## SRP043378_1
HIF1_Q3%MSIGDB_C3%HIF1_Q3;ZNF354B_TARGET_GENES%MSIGDB_C3%ZNF354B_TARGET_GENES;E12_Q6%MSIGDB_C
3%E12_Q6;COMP1_01%MSIGDB_C3%COMP1_01;TAL1BETAITF2_01%MSIGDB_C3%TAL1BETAITF2_01
## SRP044608_11
E2F1_Q3_01%MSIGDB_C3%E2F1_Q3_01;RBM34_TARGET_GENES%MSIGDB_C3%RBM34_TARGET_GENES;FOXJ2_01%MSIG
DB_C3%FOXJ2_01;WCTCNATGGY_UNKNOWN%MSIGDB_C3%WCTCNATGGY_UNKNOWN;SKIL_TARGET_GENES%MSIGDB_C3%SK
IL_TARGET_GENES
## SRP044608_2  METHYLCYTOSINE_DIOXYGENASE_TET_UNIPROT_A0A023HHK9_UNREVIEWED_TARGET_GENES%MSI
GDB_C3%METHYLCYTOSINE_DIOXYGENASE_TET_UNIPROT_A0A023HHK9_UNREVIEWED_TARGET_GENES;F10_TARGET_G
ENES%MSIGDB_C3%F10_TARGET_GENES;TTCYNRGAA_STAT5B_01%MSIGDB_C3%TTCYNRGAA_STAT5B_01;SKIL_TARGET
_GENES%MSIGDB_C3%SKIL_TARGET_GENES;ZNF391_TARGET_GENES%MSIGDB_C3%ZNF391_TARGET_GENES
## SRP043378_2
HIF1_Q3%MSIGDB_C3%HIF1_Q3;ZNF354B_TARGET_GENES%MSIGDB_C3%ZNF354B_TARGET_GENES;E12_Q6%MSIGDB_C
3%E12_Q6;COMP1_01%MSIGDB_C3%COMP1_01;TAL1BETAITF2_01%MSIGDB_C3%TAL1BETAITF2_01
## SRP043378_3
HIF1_Q3%MSIGDB_C3%HIF1_Q3;ZNF354B_TARGET_GENES%MSIGDB_C3%ZNF354B_TARGET_GENES;E12_Q6%MSIGDB_C
3%E12_Q6;COMP1_01%MSIGDB_C3%COMP1_01;TAL1BETAITF2_01%MSIGDB_C3%TAL1BETAITF2_01
## SRP063500_2
HIF1_Q3%MSIGDB_C3%HIF1_Q3;E12_Q6%MSIGDB_C3%E12_Q6;TTCYNRGAA_STAT5B_01%MSIGDB_C3%TTCYNRGAA_STA
T5B_01;MEIS1_01%MSIGDB_C3%MEIS1_01;SMTTTTGT_UNKNOWN%MSIGDB_C3%SMTTTTGT_UNKNOWN
##                          Category
## SRP043378_1  treatment + timepoint
## SRP044608_11 treatment + timepoint
## SRP044608_2  treatment + timepoint
## SRP043378_2  treatment + timepoint
## SRP043378_3  treatment + timepoint
## SRP063500_2              treatment
```

### Plotting outputs

#### Enrichment plots.

The `proccess_and_plot_DEET_enrich()` takes the output of DEET directly and makes barplots and dotplots of the most enriched comparisons at the DE, BP, and TF levels. A list of ggplot objects are returned for further manipulation.

The `DEET_enrichment_plot()` generates an invidiual barplot or dotplot to allow for increased control from the user. In this example, we will use the `proccess_and_plot_DEET_enrich()` function.

```
enr_plots <- proccess_and_plot_DEET_enrich(DEET_full_dataset,horizontal = TRUE,topn = 5, widt
h = 15)
```

```
## Removing DE_correlations element from output
```

```
## Removing non-significant DE lists
```

```
## Genarating batplot of traditional pathway enrichments
```

```
## Genarating barplot of pathway enrichment (BP + TF) if available
```

```
## Genarating barplot of DEET enrichment (DE + BP + TF) if available
```

These plots can be visualized by printing them into R, as they're the output of the ggplot objects

```
# Dotplot of enriched comparisons at DE, BP, and TF levels
print(enr_plots$DEET_DotPlot)
```

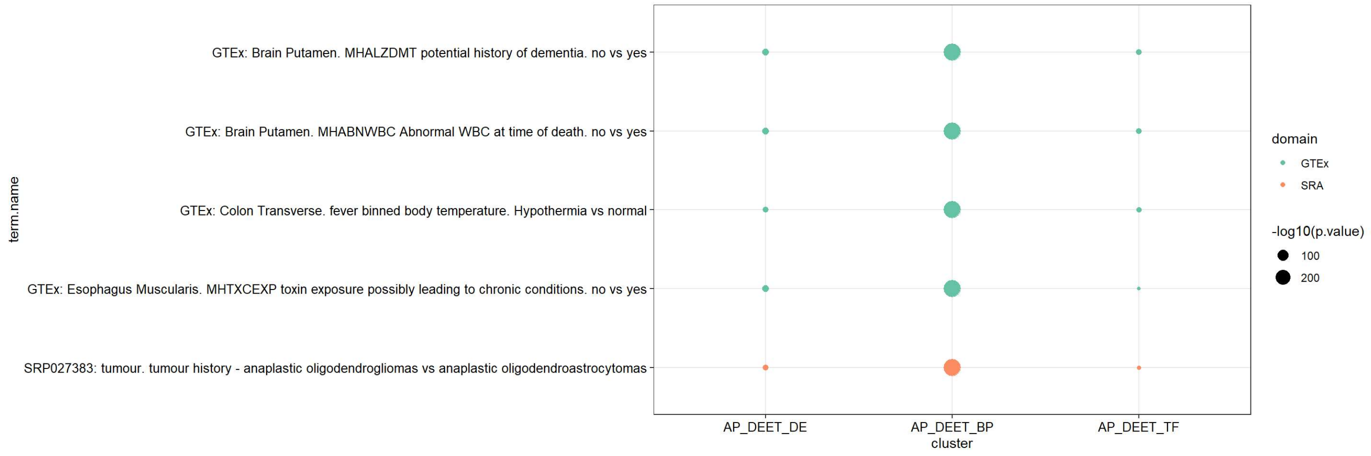


```
# Barplot of enriched comparisons at DE level
print(enr_plots$individual_barplot$AP_DEET_DE)
```

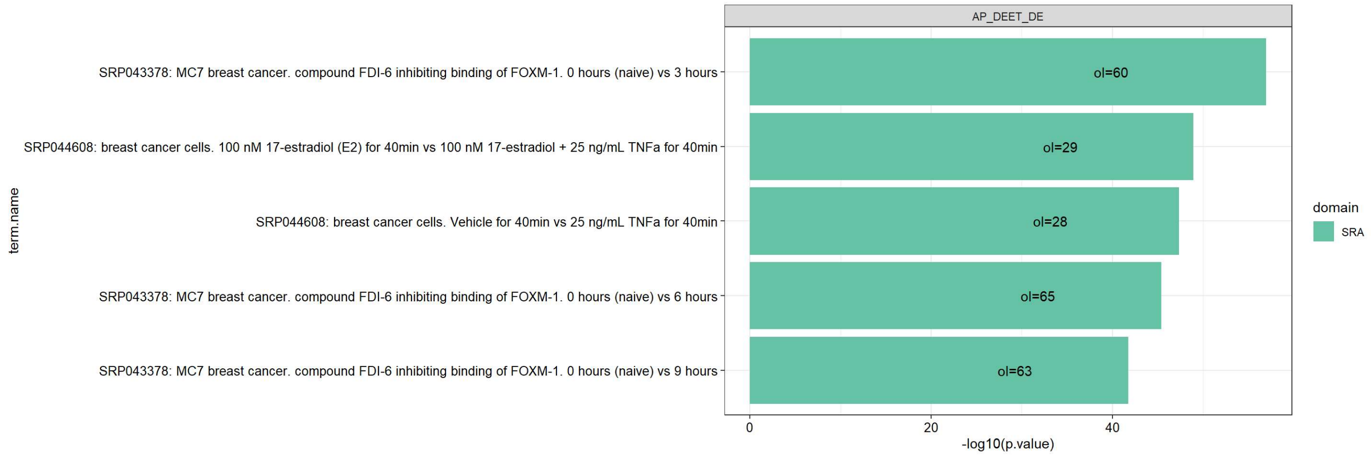


#### Correlations.

Scatterplots of indivudal enriched + correlated studies can also be plotted within DEET. The output is a list of ggplot objects containing the scatterplots of all enriched + correlated studies.

```
# Generate correlation plots

correlation_input <- DEET_full_dataset$DE_correlations
correlation_plots <- DEET_plot_correlation(correlation_input)
```

```
## There are 5 comparisons with a significant correlation.
```

```
# Print out the correlation plot of one of the studies

correlation_plots[[1]]
```

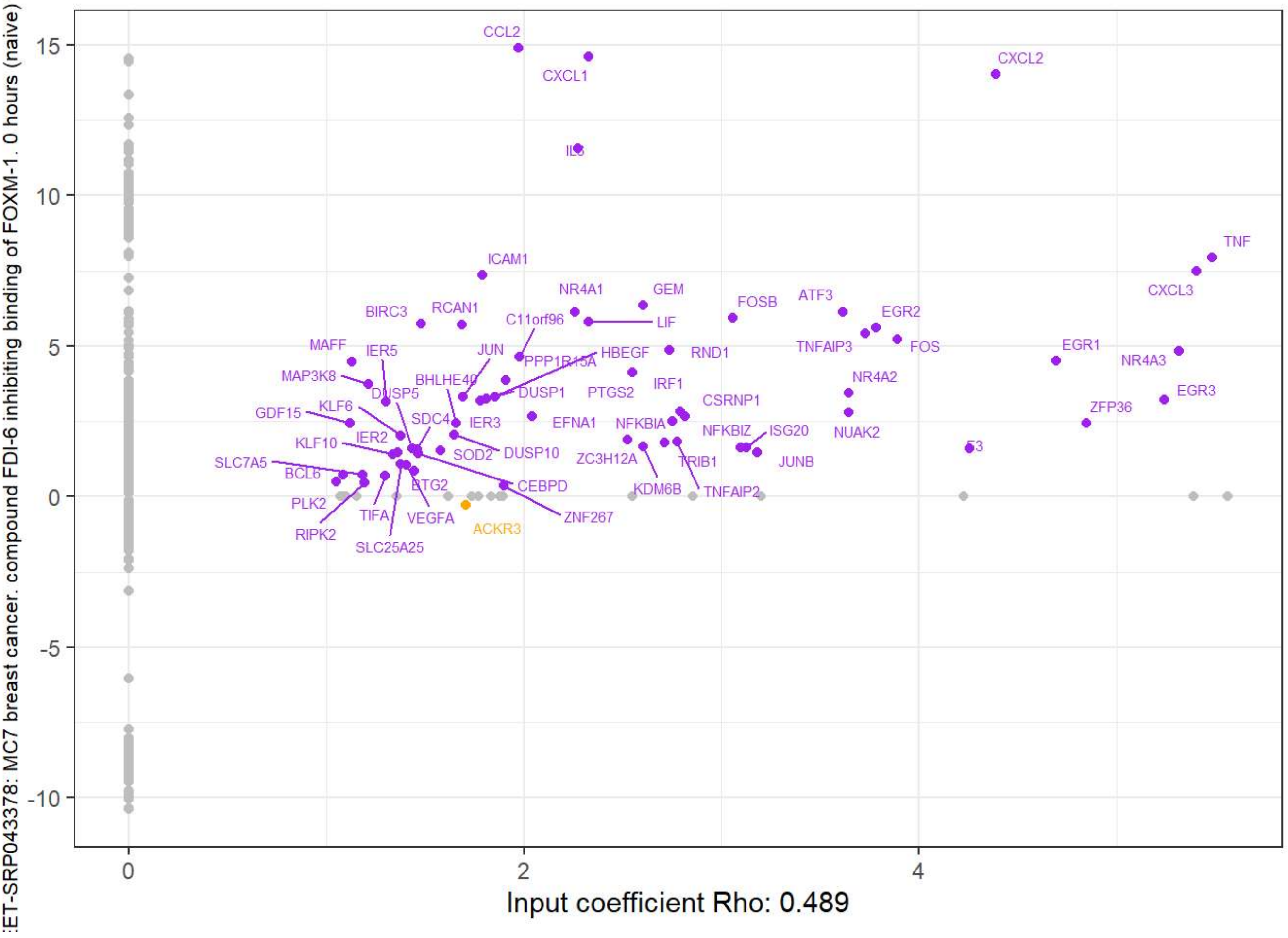


### Genes with similar DE patterns to a gene or metadata of interest.

The last built-in application of DEET is to use elastic net and correlation-based analyses to find genes with patterns of DE that represent an input variable. Simply input the response variable and the data type that the response variable is. In this example, we will identify genes whose DE expression patterns follows the pattern of the TNF gene. This is expected to that 10-20 mins on a laptop with 16Gb of RAM.

```
gene <- "TNF"
responseFC <- unlist(unname(DEET_features[gene,]))
DEET_features1 <- DEET_features[!(rownames(DEET_features) %in% gene),]

# Took the absolute value as the directionality of the "up" condition within comparisons are
 based on letters
TNF_feature_FC <- DEET::DEET_feature_extract(mat = DEET_features1, response = responseFC,data
type = "continuous")
```

With related features computed, print the output and sort on coefficient size and/or p-value to identify genes whose DE patterns best fits your input variable.

```
## Looking at the top coefficients using the elastic net

# Get coefficients
TNF_feature_FC_coef <- TNF_feature_FC$elastic_net_coefficients

# Get genes ordered by coefficient size
TNF_feature_FC_coef_enr <- names(sort(abs(TNF_feature_FC_coef), decreasing = T))

head(TNF_feature_FC_coef_enr)
```

```
## [1] "CCDC7"    "NFKBIA"   "SH3GL1P2" "SEMA4A"   "STX11"    "TCEAL1"
```

```
## Looking at top most correlated DEGs

TNF_feature_FC_cor <- TNF_feature_FC$basic_features$res

TNF_feature_FC_cor <- TNF_feature_FC_cor[order(TNF_feature_FC_cor$FDR),]
TNF_feature_FC_cor <- TNF_feature_FC_cor[TNF_feature_FC_cor$Rho > 0.2,]

head(TNF_feature_FC_cor)
```

```
##                 Rho            P           FDR
## CCL3-AS1  0.3945377 2.835740e-118 1.135799e-113
## TNFAIP3   0.3886076 1.671497e-114 6.694678e-110
## BIRC3     0.3708746 1.104523e-103  4.423726e-99
## BCL2A1    0.3700704 3.296352e-103  1.320189e-98
## LINC02605 0.3590506  7.759231e-97  3.107494e-92
## CCL3      0.3563932  2.449388e-95  9.809310e-91
```
